# Inter-protocol reproducibility of multi-shell diffusion MRI metrics across major neuroimaging study protocols

**DOI:** 10.64898/2026.11.13.766526

**Authors:** Jamie Wren-Jarvis, Valentin Stepanov, Benjamin Ades-Aron, Jenny Chen, Subah Mehrin, Nalini Jeet, Santiago Coelho, Ricardo Coronado-Leija, Dmitry S. Novikov, Jelle Veraart, Els Fieremans

## Abstract

Diffusion MRI is a core component of large-scale neuroimaging initiatives, yet the reproducibility of quantitative dMRI metrics across different acquisition protocols remains unclear. We acquired seven multi-shell diffusion protocols representative of major consortia and clinical settings on a single scanner in 41 healthy adults, isolating protocol effects from hardware and site variability. A linear mixed-effects model was used to estimate the inter-protocol coefficient of variation (CV) and the intraclass correlation coefficient (ICC) for diffusion tensor imaging (DTI), diffusion kurtosis imaging (DKI), and standard model imaging (SMI) metrics across atlas-based white matter regions. Rank agreement was assessed with Kendall’s *W*, and repeated scans of one protocol provided a reference for test-retest variability. Overall, inter-protocol variability was low, with median CVs of 2.9-6.1% for DTI, 3.7-5.0% for DKI, and 3.0-4.7% for SMI metrics. Protocols ranked subjects consistently (*W* = 0.73-0.86) but agreed less in absolute value (ICC = 0.31-0.66), with fractional anisotropy (FA) being the most stable metric. Acquisition parameters, including TE, *b*-values, and signal-to-noise ratio jointly explained 53-93% of the inter-protocol variance. Differences in the b-values used for DTI and DKI estimation were particularly important for the diffusivities: matching b-values across protocols reduced the median inter-protocol CV for MD from 5.0% to 1.8%, with smaller reductions for mean kurtosis (MK), axial kurtosis (AK) and the axonal water fraction (*f*). These findings demonstrate good reproducibility of quantitative higher-order diffusion MRI metrics across commonly used multi-shell acquisition protocols, which can be further improved by aligning *b*-values, providing a foundation for data pooling, harmonization, and normative modeling in large-scale neuroimaging studies.

## 1. Introduction

Diffusion magnetic resonance imaging (dMRI) is a noninvasive imaging technique sensitive to microstructure beyond conventional structural MRI and has become a core component of large-scale neuroimaging initiatives, including Alzheimer’s Disease Neuroimaging Initiative (ADNI), UK Biobank, Adolescent Brain Cognitive Development (ABCD), and the Human Connectome Project (HCP) (Bookheimer et al., 2019; Casey et al., 2018; Jack et al., 2008; Miller et al., 2016; Thompson et al., 2014; Van Essen et al., 2012). Multi-shell dMRI data, acquired as part of these collections, enables estimation of quantitative metrics derived from signal representations such as diffusion tensor imaging (DTI) (Basser et al., 1994) and diffusion kurtosis imaging (DKI) (Jensen et al., 2005), and multi-compartment biophysical models such as the Standard Model (Novikov et al., 2018) via standard model imaging (SMI) (Coelho et al., 2022). These metrics provide complimentary measures of white matter (WM) microstructure that can be studied across development, aging, and neurological disease (Kim et al., 2026; Liao et al., 2024; Lucignani et al., 2021; MacIver et al., 2024; Schilling et al., 2022; Villalón-Reina et al., 2026).

There is currently no standardized protocol for dMRI experiments, in part due to hardware considerations and ongoing development of analysis approaches. As a result, diffusion MRI acquisition protocols vary across studies, sites, and large-scale neuroimaging initiatives. Differences in acquisition parameters such as *b*-values, the number and distribution of diffusion directions, spatial resolution, echo and repetition time (TE/TR), and the use of acceleration techniques, may influence image quality and derived diffusion parameter estimates (Basser & Pierpaoli, 1996; Giannelli et al., 2009; Veraart et al., 2011; Vollmar et al., 2010; Zhan et al., 2010). To what extent these differences affect the reproducibility of quantitative higher-order diffusion metrics remains unclear. Establishing inter-protocol reproducibility is particularly important for large-scale neuroimaging research, where combining data across cohorts offers opportunities to increase statistical power and investigate biological effects across diverse populations.

Substantial efforts have been devoted to developing harmonization strategies to mitigate non-biological variability (Cetin Karayumak et al., 2019; Fortin et al., 2017; Jovicich et al., 2014; Mirzaalian et al., 2016; Ning et al., 2020). Methods including ComBat and RISH-based harmonization have been developed to improve cross-site compatibility by reducing batch effects (De Luca et al., 2022; Fortin et al., 2017; Hu et al., 2023). Quantifying inter-protocol variability and identifying its drivers would inform optimal protocol design for multi-site studies and advance the harmonization of dMRI data.

Previous studies have characterized the reliability of neuroimaging measures through test-retest designs (Brandmaier et al., 2018; G. Chen et al., 2018) and the variability of dMRI measures across scanners and imaging sites. Test-retest studies of advanced diffusion metrics have shown that reliability varies across metrics and brain regions and can be influenced by acquisition settings, including varying *b*-values, acceleration parameters, and image processing tools (Lucignani et al., 2021). Datasets such as MASiVar and ON-Harmony have characterized variability across scanners, vendors, and acquisition settings (Cai et al., 2021; Warrington et al., 2023, 2025). However, the specific contribution of acquisition protocol differences to measurement variability, independent of scanner related effects, remains insufficiently characterized. An inter-subject comparison on a single scanner provides a controlled way to quantify protocol related variability and evaluate it relative to test-retest variability.

In this study, we created a benchmark dataset of multi-shell dMRI protocols used by major consortia, together with a clinical multi-shell protocol, all acquired on a single scanner. This design isolates protocol-driven variability from hardware, vendor, and software factors. For DTI, DKI, and SMI metrics in white-matter regions, we quantified inter-protocol variability and reliability, assessed whether participant rankings were preserved across protocols, and tested whether inter-protocol variability exceeds test-retest variability (H1). We then examined which acquisition parameters are associated with inter-protocol variability and tested whether matching the *b*-range across protocols reduces it (H2). Our goal was to establish how reproducible dMRI metrics are across commonly used protocols and to identify practical ways to improve their comparability for large-scale neuroimaging research.

## 2. Method

### 2.1 Data Acquisition and Parameters

Forty one volunteers (21 M / 20 F; mean age 42.98 ± 18.00 years; range 23-80 years) underwent imaging on a Siemens Prisma 3T MRI scanner (**Appendix Figure A1**). All participants reported no history of neurological disorder, and no clinically significant abnormalities were identified on the research MRI examinations upon review by a board-certified radiologist. The study was approved by the local Institutional Review Board, and informed consent was obtained and documented from all volunteers prior to scanning. Each scan included a set of seven diffusion protocols performed during a single visit, listed in **Table 1**. Protocols include those defined for the Human Connectome Project on Aging and Development (HCP-DA; Bookheimer et al., 2019; Harms et al., 2018; Somerville et al., 2018), ABCD (Casey et al., 2018), ADNI (Jack et al., 2008) the UK Biobank (Miller et al., 2016), NYU’s Alzheimer’s Disease Research Center (*ADRC*), and two NYU Langone Health clinically used protocols referred to as MESO. A repeat scan of MESO with an echo time (TE) of 96 ms (MESO-96), was acquired to quantify test-retest variability. For HCP-DA, we used one set of diffusion acquisitions with opposite phase-encoding directions (HCP-DA1). We also collected a second pair using the same acquisition parameters as HCP-DA1, except for a modified *b*-value (HCP-DA2b). The NYULH protocols were used to scan approximately 8000 clinical patients and research volunteers during 2014-2020 (J. Chen et al., 2023, 2024). The protocols were implemented using either the Siemens product diffusion sequence (ABCD, ADNI, ADRC, and MESO) or the CMRR multiband diffusion sequence (Auerbach et al., 2013) (HCP-DA and UK Biobank), reproducing the original acquisition parameters as closely as possible. Similarly, both 20-channel and 32-channel head coils are used for RF signal reception, depending on the dMRI protocol specifications. For each subject, all MRI data was collected within a single scan session, but subjects were re-positioned in the scanner after changing the coils.

**Table 1.** Acquisition parameters across protocols. Seven diffusion MRI protocols were included in this study: the protocol used in Human Connectome Project on Aging and Development (HCP-DA), Adolescent Brain Cognitive Development (ABCD) study, Alzheimer’s Disease Neuroimaging Initiative (ADNI), UK Biobank, NYU Alzheimer’s Disease Research Center (ADRC), and two NYU Langone Health protocols acquired with different echo times, referred to as MESO-72 and MESO-96. The MESO-96 protocol was acquired twice per subject to enable assessment of test-retest variability. HCP-DA1 denotes the first pair of HCP Development/Aging diffusion acquisitions with opposite phase-encoding directions, whereas HCP-DA2b refers to the second pair, acquired using the same parameters as HCP-DA1 except for a modified *b*-value. For the HCP-DA1 and HCP-DA2b protocols, the reported number of *b*-value volumes [n] include acquisitions in both forward and reverse phase encoding directions. Signal-to-noise ratio (SNR) was computed within a white matter mask and averaged across subjects for each protocol.

| Protocol | TE (ms) | TR (ms) | Accel. Factor PE (iPAT) | Accel. Factor Slice | Total Accel. Factor | Partial Fourier | Coil | Total Scan Time (HH:MM:SS) | Voxel Size (mm) | FOV (mm) | Slices | Matrix size | *b*-values (s/mm<sup>2</sup>) [n] | SNR |
|---|---|---|---|---|---|---|---|---|---|---|---|---|---|---|
| **MESO-96** | 96 | 3700 | 2 | 2 | 4 | 6/8 | 20 CH | 0:06:22 | 1.7x1.7x3.0 | 220 | 50 | 130x130x50 | 0 [4], 250 [4],1000 [20], 2000 [60] | 17.92 |
| **UK Biobank** | 92 | 3600 | 1 | 3 | 3 | 6/8 | 32 CH | 0:07:08 | 2.0x2.0x2.0 | 210 | 72 | 104x104x72 | 0 [5], 1000 [50], 2000 [50] | 27.80 |
| **ABCD** | 89 | 4200 | 1 | 3 | 3 | 6/8 | 32 CH | 0:07:50 | 1.7x1.7x1.7 | 240 | 81 | 140x140x81 | 0 [7], 500 [6], 1000 [15], 2000 [15], 3000 [60] | 21.74 |
| **MESO-72** | 72 | 3700 | 2 | 2 | 4 | 6/8 | 20 CH | 0:06:14 | 1.7x1.7x3.0 | 220 | 50 | 130x130x50 | 0 [4], 250 [4],1000 [20], 2000 [60] | 24.36 |
| **ADRC** | 71 | 4000 | 2 | 2 | 4 | Off | 20 CH | 0:06:47 | 2.5x2.5x2.5 | 220 | 62 | 88x88x62 | 0 [4], 250 [4], 1000 [20], 2000 [60] | 29.46 |
| **ADNI** | 71 | 3400 | 1 | 3 | 3 | 6/8 | 20 CH | 0:08:02 | 2.0x2.0x2.0 | 232 | 81 | 116x116x81 | 0 [13], 500 [6], 1000 [48], 2000 [60] | 23.94 |
| **HCP-DA1** | 89 | 3230 | 1 | 4 | 4 | 6/8 | 32 CH | 0:11:22 | 1.5x1.5x1.5 | 210 | 92 | 140x140x92 | 0 [7], 1500 [50], 3000 [50] | 16.38 |
| **HCP-DA2b** | 89 | 3230 | 1 | 4 | 4 | 6/8 | 32 CH | 0:11:14 | 1.5x1.5x1.5 | 210 | 92 | 140x140x92 | 0 [7], 1000 [49], 2000 [50] | 16.42 |

### 2.2 Image Processing

All dMRI data were preprocessed using the DESIGNER pipeline (J. Chen et al., 2024). The pipeline utilized MPPCA denoising (Cordero-Grande et al., 2019; Veraart et al., 2016) to reduce thermal noise. Gibbs correction was applied using a sub-voxel shift approach that accounts for Partial Fourier encoding direction and factors (H. Lee et al., 2021). In absence of Partial Fourier encoding, the original sub-voxel shift approach of (Kellner et al., 2016) was applied. Thereafter, subject motion and geometric distortions from susceptibility and eddy current effects were corrected using FSL’s Topup and Eddy (Andersson et al., 2003; Smith et al., 2004). A b = 0 normalization was applied, in which all diffusion-weighted images were rescaled using smoothed (3 mm Gaussian kernel) b = 0 images to correct voxelwise intensity differences across series. For HCP-DA data, denoising was independently applied to imaging series with reversed phase encoding.

### 2.3 dMRI parametric maps

After preprocessing, the following parametric maps were computed using DESIGNER’s Tissue Microstructure Imaging toolbox (documentation available at https://nyu-diffusionmri.github.io/DESIGNER-v2/docs/TMI: mean diffusivity (MD), radial diffusivity (RD), axial diffusivity (AD), and fractional anisotropy (FA) using DTI, mean kurtosis (MK), radial kurtosis (RK), and axial kurtosis (AK) using DKI with fast estimation using the angular average of the cumulant (Ades-Aron et al., 2025; Hansen et al., 2013), and axonal water fraction (*f*) and fiber dispersion (*p*_2_) using SMI. We did not report the SMI compartmental diffusivities because, for the available two-shell protocols with b ≤ 2000 s/mm² and a single TE, their estimates may be strongly regularized by the data-driven estimator, potentially obscuring subtle differences between experimental conditions (Coelho et al., 2024). In the main analysis, DTI parameteric maps were computed using *b*-values up to 1500 s/mm^2^ and DKI and SMI maps were computed with the entire *b-*range of each protocol. To see how parametric maps compare when *b*-values are matched, an additional analysis was done using the same *b*-values across protocols, i.e. *b*=0, and *b*=1000 s/mm^2^ for DTI and *b*=0, *b*=1000 and *b*=2000 s/mm^2^ for DKI and SMI parametric maps. In both computational versions, windowed adaptive nonlocal means smoothing (Wiest-Daesslé et al., 2007) and diffusion-*b*0restore (J. Chen et al., 2026) was applied to the diffusion weighted images prior to DKI model estimation.

### 2.4 Image Registrations and ROI Data Extraction

Diffeomorphic image registration between individual dMRI data and the JHU-ICBM-DTI-81 atlas template was applied using Advanced Normalization Tools (Tustison et al., 2021) (ANTs) to provide an automated segmentation of the WM in the 48 widely-adopted regions of interest (ROI). Total WM was defined as the union of JHU-ICBM ROIs. The FA maps of the study data and the ICBM-152 atlas template were used as the moving and fixed image, respectively. The inverse affine transformation matrix and warp field were applied to transform the 48 JHU-ICBM ROI labels into the native space of each subject specific protocol, using GenericLabel to maintain the label structure of the atlas. Average values from each diffusion parameter map of each subject were extracted from within each JHU-ICBM ROI and WM mask and used in further analyses. Within each ROI, for each metric, voxelwise outliers were excluded prior to computing summary statistics: Voxels overlapping cerebrospinal fluid were removed using a mask obtained by thresholding the mean-diffusivity map (MD ≤ 2.5 μm²/ms), and only voxels with non-negative, finite values were kept. The mean and standard deviation of the remaining voxels were computed, and voxels more than five standard deviations from the regional mean were removed. Regional statistics were then computed on the surviving voxels. This ROI extraction was done for each region, metric, protocol and subject; after filter application 27 ROIs for different metrics were excluded because no valid voxels remained (**see Supplementary Figure S2**).

### 2.5 SNR Computation

Average SNR from within WM and individual ROIs was computed for each dMRI protocol. For each subject and protocol, SNR was calculated on a voxel-wise basis as the ratio of the non-DWI (b=0) signal intensity to the spatially varying noise map, and averaged within respective ROIs. Subject-averaged WM SNR values are computed.

### 2.6 Repeatability and reproducibility assessments

Repeatability and reproducibility were evaluated across repeated scans, subjects, and acquisition protocols. Variability was summarized using coefficients of variation (CVs), absolute agreement using the intraclass correlation coefficient (ICC), and rank-order agreement using Spearman’s ρ and Kendall’s *W*. Analyses were performed separately for each diffusion metric and ROI. **Appendix Figure A2** summarizes the data included in each analysis. Detailed model specifications, equations, and computational procedures are provided in the Supplementary Methods.

#### 2.6.1 Model-based variability and absolute agreement

Linear mixed-effects models (LME) (Laird & Ware, 1982) were fitted for each metric and ROI using restricted maximum likelihood in R v4.5.2 (lme4 v2.0-1). Protocol and age were included as fixed effects, and subject as a random intercept. The models estimated inter-subject and residual variances, along with an age-adjusted mean for each protocol (Bernal-Rusiel et al., 2013). Inter-protocol variance was the variance of these protocol means. The ICC was computed using the two-way, fixed-rater ICC for single measurements, ICC(A,1) (Shrout & Fleiss, 1979), derived as the inter-subject variance divided by the sum of the inter-subject, inter-protocol, and residual variances. This measure treats systematic differences between protocols as disagreement. All CVs were expressed relative to a common reference mean averaged over the full and matched *b*-value analyses (Methods 2.8.1), so that all CVs share the same scale. The inter-protocol CV (*CV_protocol_*) was the square root of the inter-protocol variance relative to this reference.

#### 2.6.2 Within-subject variability

Measurement noise was estimated from subjects for whom both the test and retest MESO-96 scans passed QC (n = 33). The test-retest CV (*CV_ws_*) was the root-mean-square of the paired differences divided by √2, relative to the common reference mean.

#### 2.6.3 Pairwise and rank-order agreement

For each pair of protocols, we calculated the age-adjusted percentage difference, the within-subject CV between protocol pairs (*CV_ws_*_, *pair*_), and Spearman’s ρ. *CV_ws_*_, *pair*_ captures both systematic protocol differences and residual within-subject variability, whereas Spearman’s ρ assesses whether protocols preserve subject rankings. Rank-order agreement across all protocols was assessed using Kendall’s coefficient of concordance, *W* (Kendall & Smith, 1939), among subjects with data from every protocol. Kendall’s *W,* unlike the ICC, is unaffected by systematic differences between protocols.

### 2.7 Cumulative contribution of protocols to inter-protocol variability

To understand how inter-protocol variability accumulates as acquisition protocols are pooled into a combined dataset, we performed a compounding analysis. Protocols were added one at a time, starting from a pair. For each subset, *CV_protocol_* was recomputed from that subset’s fitted protocol effects (from the LME in 2.6.1), relative to the pooled intercept. All 2,520 unique orderings were evaluated. Each was scored by the Frobenius norm across the nine metrics and six compounding steps, where the lowest norm was considered optimal. The search was repeated within each metric family (DTI, DKI, SMI) and with the three families weighted equally. The analysis was run in whole white matter and repeated in the 27 hemisphere-averaged regions, with uncertainty quantified by a 200-replicate subject bootstrap in which the model was refit on each replicate (**Supplementary Figure S6**).

### 2.8 Contribution of scan parameters to inter-protocol variability

We tested whether inter-protocol differences could be attributed to three acquisition parameters: TE, maximum *b*-value, and WM signal-to-noise ratio (SNR). This analysis used all eight protocols (**Appendix Figure A2**) in the 27 bilateral regions. For DTI metrics, the maximum *b*-value was the highest *b*-value used for DTI estimation (*b*=1000 or *b*=1500 s/mm^2^); for DKI and SMI metrics, this value and the highest *b-*value in the *b-*range (*b*=2000 or *b*=3000 s/mm^2^) were included. The LME was refit with protocol replaced by these parameters as fixed effects, retaining age and the subject random intercept. Each coefficient was divided by the standard deviation (SD) of the metric in the region, expressed in metric SDs per protocol-level SD of the parameter, and reported as medians and interquartile ranges across regions. The eight fitted protocol effects were then regressed on TE, SNR, and the maximum *b*-value matching the metric’s estimation range by ordinary least squares. *R*² from this regression gives the fraction of inter-protocol variance explained by the parameters jointly, and is reported with adjusted *R*², which corrects for the number of parameters relative to the number of protocols. Variance inflation factors were computed for each model.

### 2.9 Statistical tests

Two prespecified hypotheses were tested for each metric using the 27 bilateral regions as paired observations: *CV_protocol_* will exceed *CV_ws_* (H1), and *CV_protocol_* will be lower when metrics are computed with matched *b-*values (*b*=0,1000, and 2000 s/mm^2^) than computed from each protocol’s full *b*-range (H2). One-sided Wilcoxon signed-rank tests were used, and effect sizes were reported using the matched-pairs rank-biserial correlation. For each hypothesis, the nine metric-level tests formed a separate family and were Bonferroni-corrected, with *p*_corr_ < 0.05 considered statistically significant. All other analyses were descriptive.

## 3. Results

### 3.1 Participants and analytic sample

Forty-one participants were scanned in a single session, yielding 332 diffusion datasets (**Appendix Figure A3**). HCP-DA2b was introduced partway through the study and acquired for the final 16 participants. Among the other eight acquisitions, 12 datasets were not acquired across six participants: six MESO TE = 96 Run 2, three ADRC, and one each from ABCD, HCP-DA1, and UK Biobank. Seventeen datasets were excluded: 16 by FSL SQUAD quality control (Bastiani et al., 2019) because of excessive motion or outlier slices and one because of an acquisition-parameter deviation. Visual quality control of the parameter maps and registrations identified no additional exclusions. Exclusions were distributed across protocols (**Appendix Table A1**). Missingness and exclusions were handled at the participant-protocol level, and all available observations were included in the mixed-effects models. The primary LME analysis included 299 datasets from all 41 participants. The remaining 16 datasets that passed quality control were from HCP-DA2b and were included only in the acquisition-parameter decomposition and matched *b*-value comparison.

Sample sizes varied by protocol and analysis because not all participants completed every protocol or passed quality control. Protocol-specific sample sizes ranged from 36 participants for ADRC to 41 for MESO TE = 96, with both runs combined. Thirty-three participants had two MESO TE = 96 acquisitions that passed quality control and comprised the test–retest sample. Twenty-eight participants had data from all seven protocols required for the Kendall’s *W* analysis. At the regional level, 0.04% of values were unavailable.

### 3.2 Inter-protocol variability and agreement

In **Figure 1**, we show *CV_protocol_* (A: full *b*-range and B: selected *b*-range) and *CV_ws_* (C) for all diffusion metrics and brain regions. On average, the *CV_protocol_* is 3.00-6.54%, 3.79-4.73%, and 3.16-4.99% for DTI, DKI, and SMI metrics, respectively. In comparison, *CV_ws_* is 1.38-3.76%, on average. *CV_protocol_* exceeds *CV_ws_* significantly for all metrics, with exception of RK. We test the robustness of these findings to outliers (**Supplementary Table S2**).

**Figure 1.**
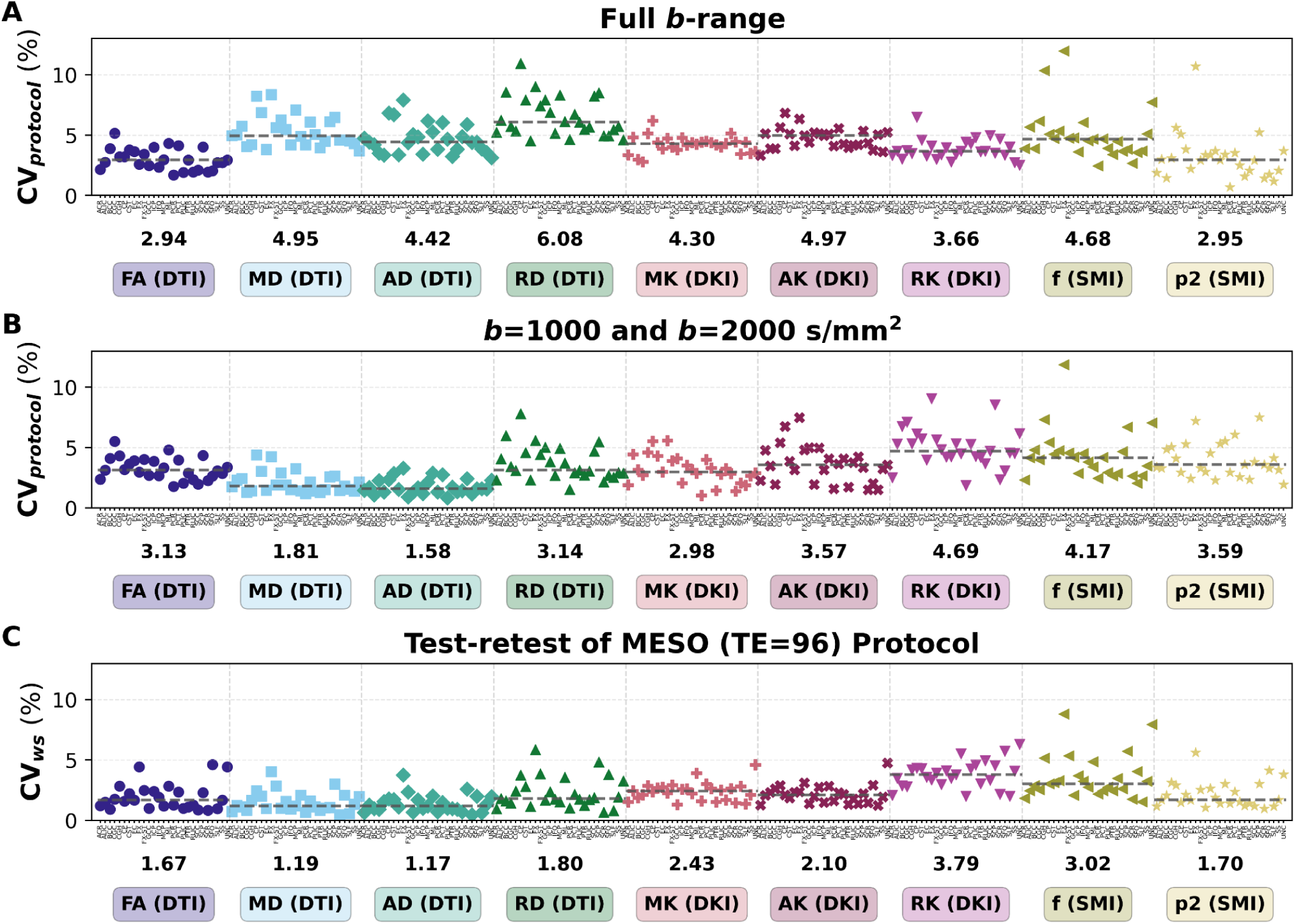
Variation between protocols with the full and matched b-value ranges, compared with Test-retest variation, for nine diffusion metrics in each white matter region. **(A)** Inter-protocol coefficient of variation *CV_protocol_*, the spread of the seven protocol means relative to the intercept (grand mean), with each protocol’s full *b*-range. **(B)** *CV_protocol_* after restricting every protocol to matched *b*-values: *b* = 0 and 1000 s/mm² for DTI, and *b* = 0, 1000 and 2000 s/mm² for DKI and SMI, with HCP-DA2b in place of HCP-DA1. **(C)** Test-retest within-subject coefficient of variation *CV_ws_* from the two MESO-96 runs (n = 33), full *b*-range. All three panels use the same denominator, the mean of the full and matched *b*-range intercepts, so they can be compared directly. Each point is one region, with values averaged across hemispheres. The whole WM mask is excluded. Dashed lines and the numbers below each group give the median across regions

We map the *CV_protocol_* across white matter regions for all diffusion metrics in **Figure 2**. *CV_protocol_* varies spatially, with the fornix (FX) and brainstem tracts (CP, SCP and CST) having the largest *CV_protocol_* In contrast, the posterior corona radiata (PCR), superior fronto-occipital fasciculus (SFO) and the posterior thalamic radiation (PTR) have the smallest *CV_protocol_* . Regional *CV_protocol_* was also only weakly related to region size (**Supplementary Figure S3**).

**Figure 2.**
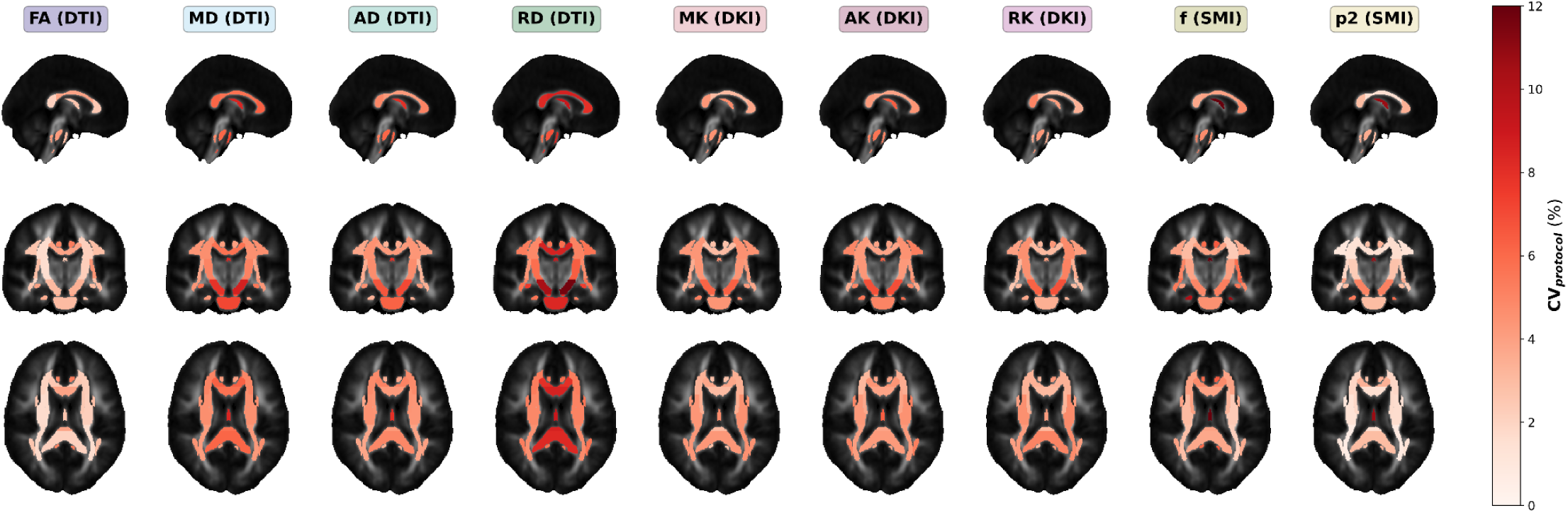
*Inter-protocol variation mapped onto the JHU white matter atlas*. Each JHU region is colored by its inter-protocol variation (*CV_protocol_*) for each diffusion metric (columns), shown on mid-sagittal (top), mid-coronal (middle) and mid-axial (bottom) slices. Left and right hemispheres keep their own values. All panels share the 0-12% color scale; darker red means more variation between protocols.

In **Figure 3**, we show Kendall’s W (left) and ICC (right) for all diffusion metrics to quantify rank order agreement and absolute agreement across protocols. The absolute agreement was poor to moderate, with ICC values ranging from 0.31 to 0.66 (**Figure 3**). In contrast, rank order agreement across the same protocols was high: Kendall’s *W* ranged from 0.73 (AK) to 0.86 (FA) (**Figure 3)**. The metrics with the lowest ICC therefore still ranked subjects consistently, while differing substantially in absolute value between protocols. Regional maps of Kendall’s *W* and ICC are shown in **Supplementary Figure S4**.

**Figure 3.**
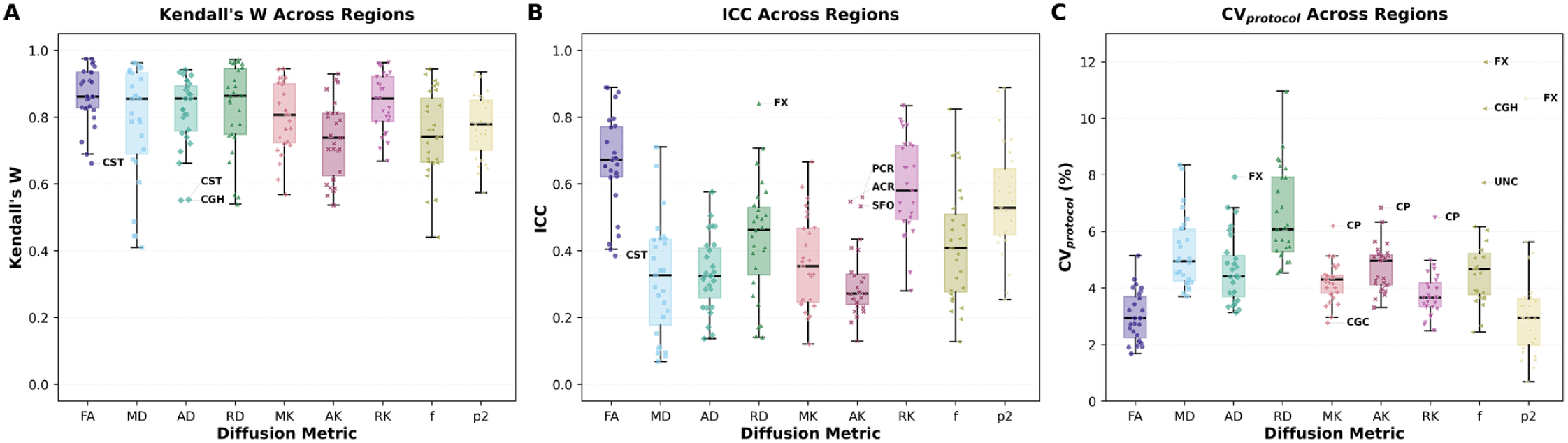
*Consistency of subject ranking and absolute agreement across protocols, by diffusion metric*. **(A)** Kendall’s W: whether the seven protocols rank subjects in the same order. **(B)** Intraclass correlation coefficient (ICC): the share of total variance that comes from differences between subjects rather than protocols. **(C)** Inter-protocol coefficient of variation (*CV_protocol_*) with each protocol’s full b-range. Each point is one white matter region, with values averaged across hemispheres. Boxes show the median and interquartile range, and whiskers extend to 1.5 × IQR. Labelled regions lie outside the whiskers. For *W* and ICC measures, 1 is perfect agreement.

### 3.3 Pairwise protocol comparisons

Inter-protocol variability and correlation between each pair of protocols are quantified by *CV_ws_*_,*pair*_ and Spearman’s ρ, respectively, shown in **Figure 4**. Disagreement was concentrated in pairs including HCP-DA1 for the diffusivities and in pairs including ABCD for the kurtosis metrics. For MD, AD and RD, all six pairs including HCP-DA1 had a *CV_ws_*_, *pair*_ of 5.83% or more. Summarized as the median across the six pairs that include each protocol, HCP-DA1 had medians of 8.31%, 6.70% and 10.66%, respectively. HCP-DA1 also had the highest median for f (5.73%), and no other protocol exceeded

**Figure 4.**
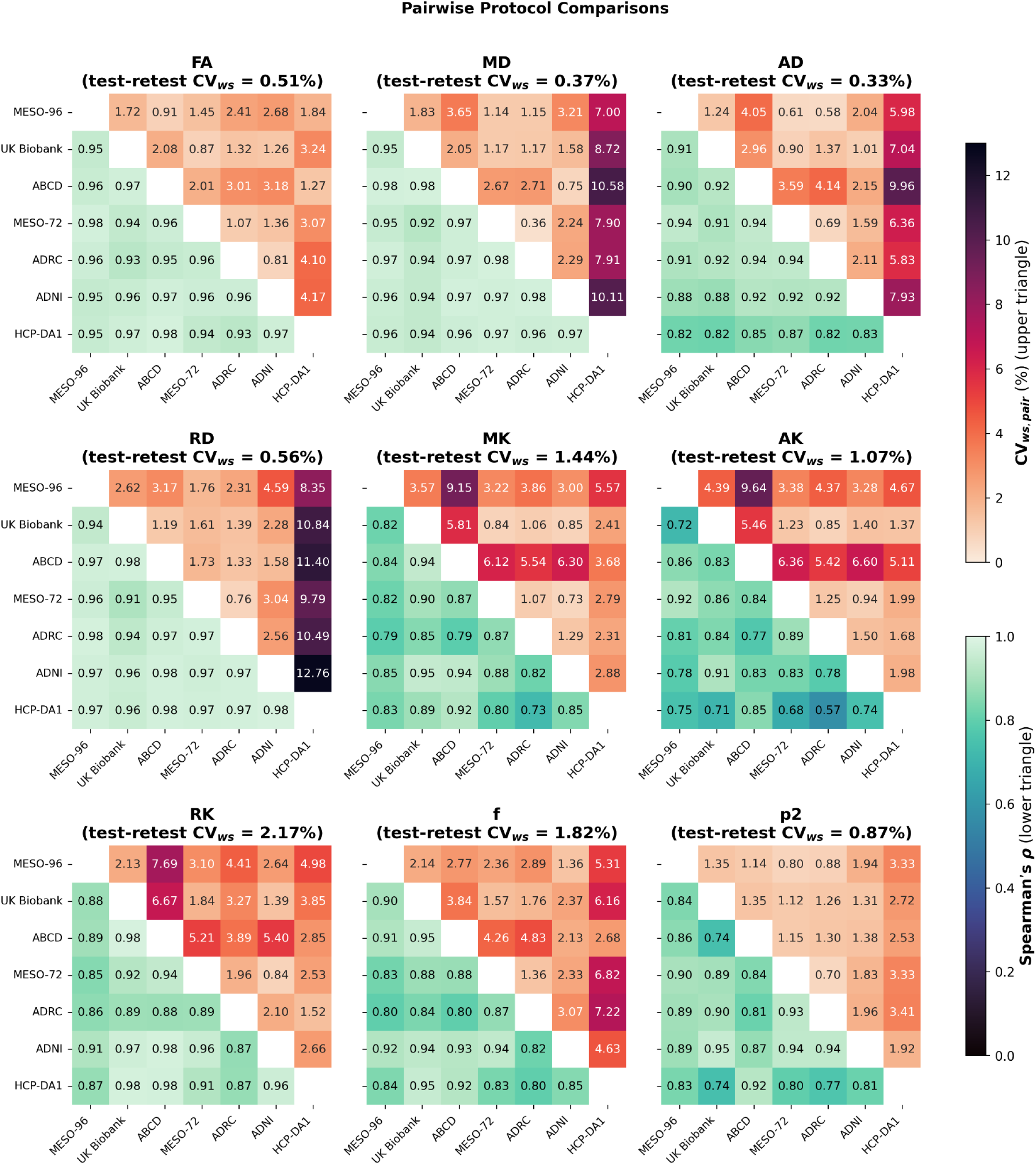
*Within-subject variation between each pair of protocols in total white matter, compared with test-retest variation*. Each panel is one diffusion metric. **Upper triangle**: pairwise within-subject coefficient of variation (*CV_ws_*_, *pair*_) for each pair of protocols. It uses the same formula and denominator (the intercept from the LME) as test-retest *CV_ws_*, so the two can be compared directly. **Lower triangle**: Spearman’s ρ between the two protocols across subjects, which shows whether they rank subjects in the same order. Each pair uses the subjects scanned with both protocols (n = 33-38). MESO-96 uses run 1 only, so every protocol contributes a single scan. The test-retest *CV_ws_* in each panel title, from the two MESO-96 runs, is a proxy for test-retest noise floor: pairs well above it differ by more than repeat-scan noise. Darker upper cells mean more variation between protocols, and darker lower cells mean less consistent subject ranking.

3.31%. For MK, AK and RK, ABCD had the highest medians (5.97%, 5.91% and 5.31%, respectively), and no other protocol exceeded 4.38%. Spearman’s ρ had medians of 0.83 (AK) to 0.97 (MD and RD), so even the pairs with the largest *CV_ws_*_,*pair*_ ranked subjects consistently. AK in pairs with HCP-DA1 showed the clearest loss of rank agreement (ρ = 0.57-0.71). Protocol variability is primarily reflected as systematic shifts in nominal values of the metrics. **Supplementary Figure S5** shows relative differences in mean value between each pair of protocols. Despite these differences in metric values, the protocols ordered subjects similarly.

### 3.4 Protocol compounding order

In **Figure 5**, we show the optimal ordering of protocols to pool data cumulatively while minimizing the inter-protocol variability. An exhaustive search over all 2,520 unique sequences identified ADRC, MESO-72, UK Biobank, ADNI, MESO-96, ABCD, HCP-DA1 as the best performing whole WM ordering across all nine metrics. The same sequence remained optimal when the three families were weighted equally, and the DTI and SMI families with only ADNI and MESO-96 exchanged. The best ordering for the DKI family was ADNI, MESO-72, UK Biobank, ADRC, HCP-DA1, MESO-96, ABCD. In each case, the protocol placed last was the one with the lowest pairwise agreement for that family: HCP-DA1 for DTI and SMI, and ABCD for DKI. This held regionally. HCP-DA1 was last in the optimal all metric ordering of all 27 regions, although only 4 regions reproduced the whole WM ordering exactly (**Supplementary Figure S6**). Along the optimal ordering, variability remained low until the last two protocols were added. The combined *CV_protocol_* across the nine metrics rose from 2.21% for the first pair to 5.27% after five protocols. It then increased by 3.21 points when ABCD was added and by 3.44 points when HCP-DA1 was added, reaching 11.91% (**Figure 5A**). The two steps affected different families (**Figure 5B**) Adding HCP-DA1 at the final step roughly doubled *CV_protocol_* for MD (1.95% to 4.98%) and AD (2.20% to 4.33%), and almost tripled it for RD (2.10% to 5.96%). The DKI metrics increased earlier, as MESO-96 and then ABCD were added (MK from 0.50% to 4.11% and AK from 0.88% to 4.33%), and declined slightly with HCP-DA1. The SMI metrics were the most stable: f rose gradually to 3.57%, and *p*_2_ remained at 1.63% or less. The protocols with the highest maximum *b*-values therefore contributed the largest increases in variability.

**Figure 5.**
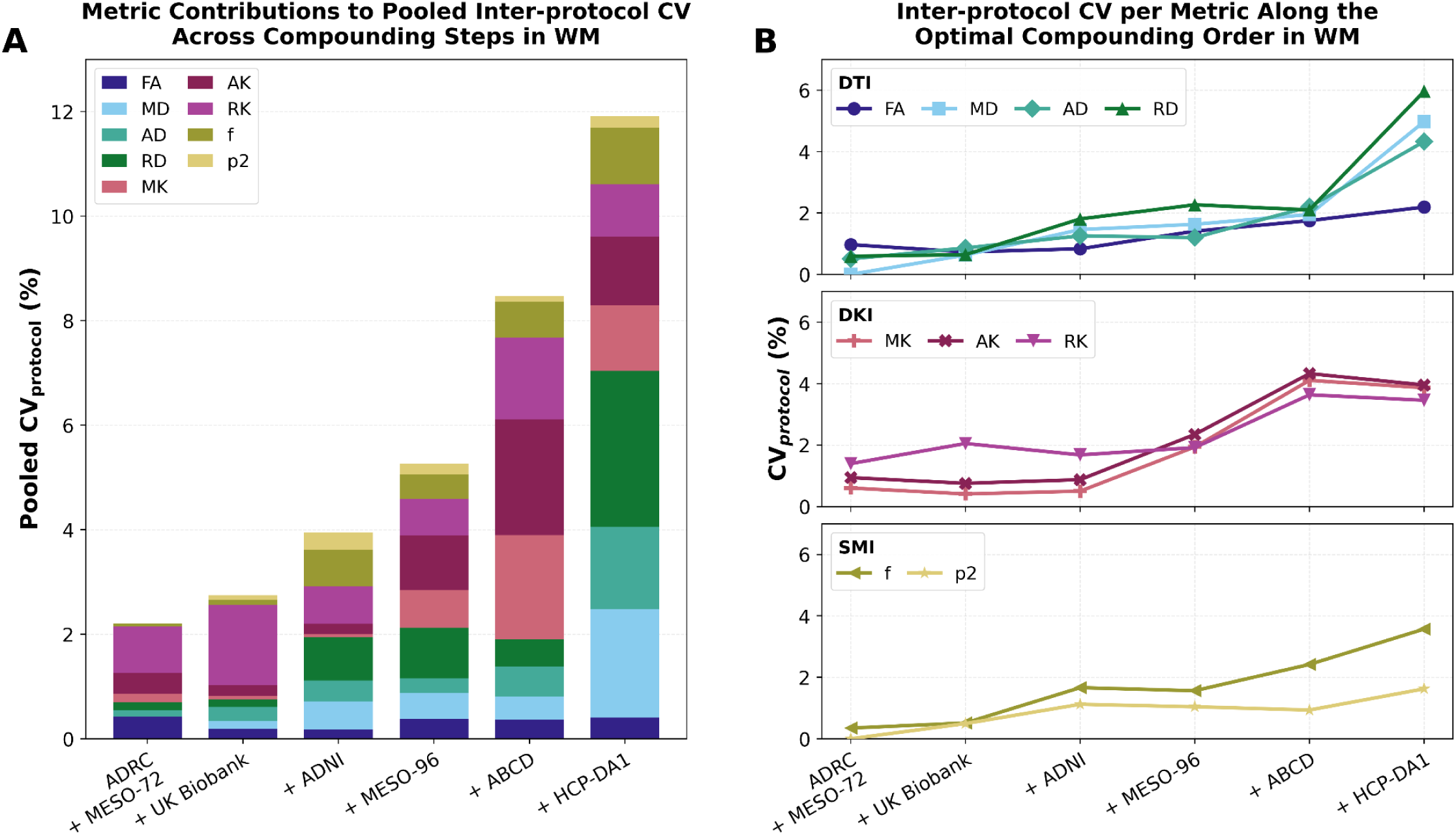
*How inter-protocol variation in total white matter accumulates as protocols are pooled.* Protocols were added one at a time in the order that kept pooled variation lowest, found by exhaustive search over all 2,520 orderings: ADRC + MESO-72, then UK Biobank, ADNI, MESO-96, ABCD and HCP-DA1. **(A)** Pooled inter-protocol coefficient of variation, √(Σ *CV_protocol_*²) over the nine metrics, at each step. Each bar is divided according to each metric’s share of Σ *CV_protocol_*². **(B)** *CV_protocol_*for each metric along the same order, grouped by model family (DTI, DKI, SMI). At every step, *CV_protocol_* is computed from age-adjusted fitted protocol effects from the mixed-effects model, is corrected for sampling noise, and is expressed relative to the pooled intercept, which is fixed across all all steps. Changes between steps therefore reflect only the change in spread between protocols, and the final step equals the full, whole study *CV_protocol_*.

### 3.5 Acquisition parameter decomposition

In Figure 6, we show the relative contribution of TE, maximum *b*-value, and WM SNR to observed variability, quantified using standardized effect size, in all diffusion metrics. These acquisitions parameters were only weakly collinear, with variance inflation factors (VIF) of 1.29 to 1.83 in the DTI model and 1.56 to 2.01 in the DKI/SMI model with both *b*-value terms. Moreover, TE, maximum *b*-value and SNR explained collectively 68.8-93.1% of the protocol variance for eight metrics, but only 52.7% for *p*₂ (**Table 2**).

**Figure 6.**
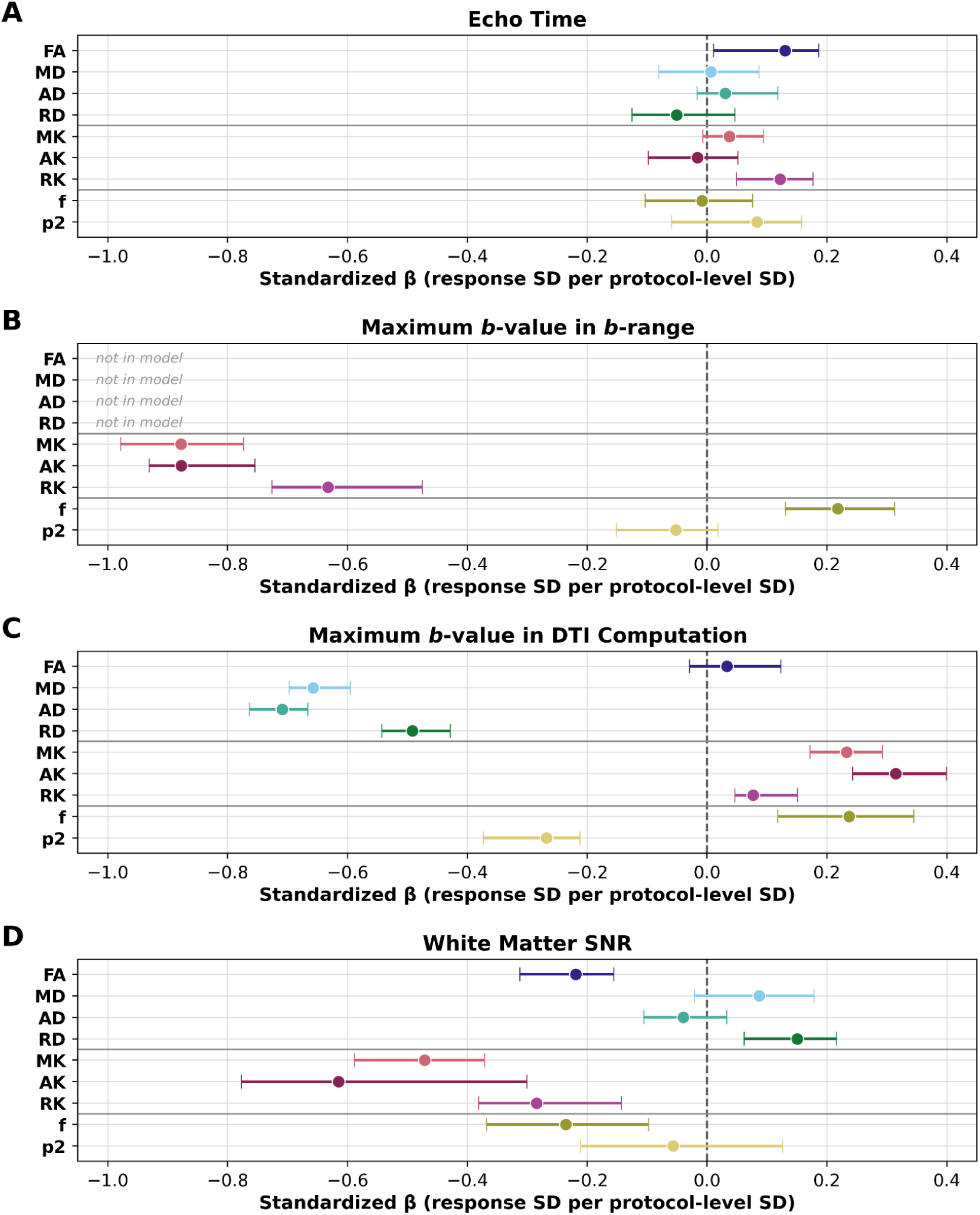
*Association of acquisition parameters with diffusion metric values*. Standardized fixed effects of **(A)** echo time, **(B)** the maximum *b*-value used for the DKI and SMI estimations, **(C)** the maximum *b*-value used for the DTI estimation and **(D)** white matter (WM) SNR. They come from mixed-effects models with the acquisition parameters as fixed effects, age as a covariate and a random intercept for each subject, fitted separately for each metric and region. The models cover eight protocols: the seven main protocols plus HCP-DA2b, which provides the only *b*-value contrast within HCP-DA1. Each parameter was z-scored across the eight protocols, so β is the change in the metric, in SDs of the metric, per SD of the parameter across protocols. DTI metrics were modelled with echo time, DTI maximum *b*-value and SNR. DKI and SMI metrics also included the DKI/SMI maximum *b*-value, so DTI metrics are “not in model” in (B). Points show the median across the 27 white matter regions (left and right hemispheres averaged), and bars show the interquartile range. The dashed line marks β = 0.

**Table 2.**
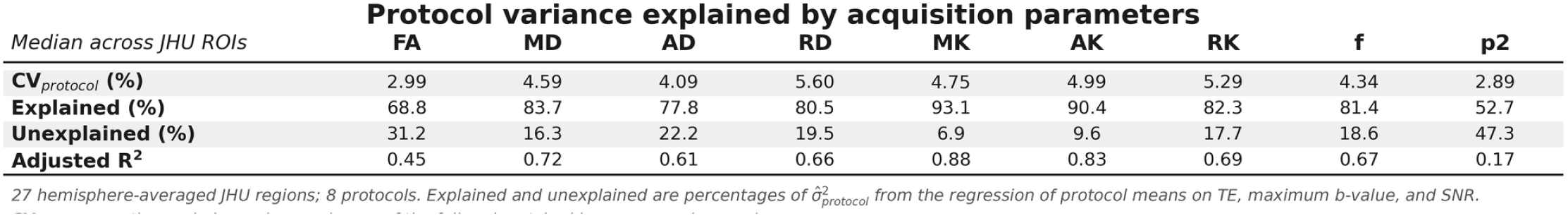
Share of protocol variance explained by TE, maximum *b*-value, and SNR (medians across 27 regions; eight protocols).

| *Median across JHU ROIs* | **FA** | **MD** | **AD** | **RD** | **MK** | **AK** | **RK** | **f** | **p2** |
|---|---|---|---|---|---|---|---|---|---|
| **CV<sub>*protocol*</sub> (%)** | 2.99 | 4.59 | 4.09 | 5.60 | 4.75 | 4.99 | 5.29 | 4.34 | 2.89 |
| **Explained (%)** | 68.8 | 83.7 | 77.8 | 80.5 | 93.1 | 90.4 | 82.3 | 81.4 | 52.7 |
| **Unexplained (%)** | 31.2 | 16.3 | 22.2 | 19.5 | 6.9 | 9.6 | 17.7 | 18.6 | 47.3 |
| **Adjusted R<sup>2</sup>** | 0.45 | 0.72 | 0.61 | 0.66 | 0.88 | 0.83 | 0.69 | 0.67 | 0.17 |

*27 hemisphere-averaged JHU regions; 8 protocols. Explained and unexplained are percentages of σ̂<sup>2</sup><sub>protocol</sub> from the regression of protocol means on TE, maximum b-value, and SNR.*
[illegible]

Maximum *b*-value had the largest and most consistent effects on diffusion metrics. The maximum *b* -value of DTI analyses had negative effects on AD (−0.71), MD (−0.66), and RD (−0.49), with the same sign in every region (**Figure 6C**). The DKI/SMI maximum *b*-value had negative effects on MK (−0.88), AK (−0.88), and RK (−0.63), also in every region (**Figure 6B**). Both *b*-values had smaller effects on the remaining metrics, including almost none on FA or *p*_2_. The kurtosis metrics depended on both shells: including the DTI maximum *b*-value in the DKI/SMI model lowered AIC in 126 of 135 metric and region cells and BIC in 118. SNR had its largest effects on the kurtosis metrics, negative for AK (−0.62), MK (−0.47), and RK (−0.28) in every region, and smaller effects elsewhere (**Figure 6D**). TE had the weakest effects of the three parameters, 0.13 or less in absolute value for every metric (**Figure 6A**).

### 3.6 Matched b-value comparison

We quantified the impact of matching *b*-values across protocol on *CV_protocol_*, see Figure 1B and Table 3. Using select *b*-values of *b=*0, 1000 and 2000 s/mm² significantly reduced *CV_protocol_* for six metrics (MD, AD, RD, MK, AK and *f*). FA, RK and *p*_2_ were not reduced. Diffusivities showed the largest reductions. *CV_protocol_* fell by about two thirds for MD and AD and by about half for RD, and matched *CV_protocol_* was lower than full *b*-range *CV_protocol_* in every region (*p_corr_*< 0.001). With matched *b*-values, *CV_protocol_* for MD and AD (medians 1.81% and 1.58%) approached test-retest *CV_ws_* (1.19% and 1.17%; **Figure 1B-C**). *CV_protocol_* fell by about 30% for MK and AK (*p_corr_* < 0.001) and by about 10% for *f* (*p_corr_* = 0.008). For FA, RK and *p*_2_, matched *CV_protocol_* increased in most regions (**Supplementary Figure S7**).

**Table 3.**
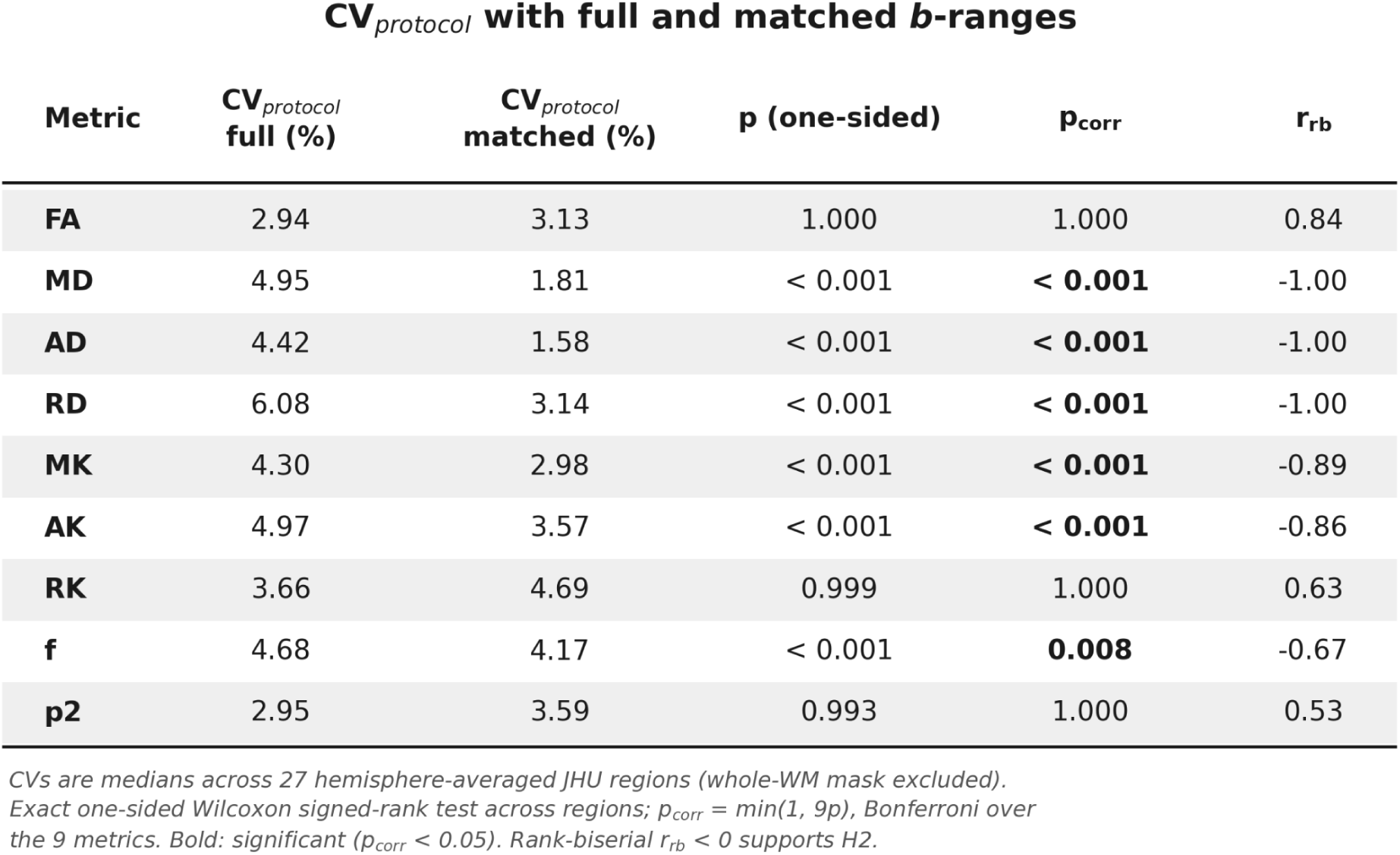
*One-sided tests of whether matching b-values reduce inter-protocol variability*. For each diffusion metric, *CV_protocol_* computed with the full and matched *b*-value ranges was compared across 27 JHU regions (hemispheres averaged, whole WM mask excluded) with an exact one-sided Wilcoxon signed-rank test of whether matching reduces it. Both CVs share a denominator, the mean of the two analyses’ intercepts. CV values are medians across regions. *p*_corr_ is Bonferroni-corrected over the nine metrics, and bold marks *p*_corr_ < 0.05. The *r*_rb_ is the matched pairs rank-biserial correlation, an effect size for how consistently regions differ in one direction. An *r* < 0 indicates lower *CV_protocol_*with matched *b*-values, and *r* = −1 means every region differs in that direction.

## 4. Discussion

Across seven multi-shell dMRI protocols, adapted from large-scale neuroimaging studies (Bookheimer et al., 2019; Harms et al., 2018; Somerville et al., 2018; Casey et al., 2018; Jack et al., 2008; Miller et al., 2016), acquired on a single scanner, inter-protocol variability in DTI, DKI, and SMI metrics ranged from about 3% to 7% (**Figure 1**), and protocols largely preserved the ranking of participants (**Figure 3-4**). Matching *b*-values across protocols reduced variability in MD, AD, RD, MK, AK, and *f* to less than 5% (**Figure 1**). Among the acquisition parameters examined, maximum *b*-value was more strongly associated with inter-protocol variability than TE or WM SNR for the DTI and DKI metrics (**Figure 6**). Most of the variability came from the two protocols that acquired *b* = 3000 s/mm², whereas the other five reached at most 2000 s/mm². HCP-DA1, which was also the only protocol with a 1500 s/mm² DTI shell, accounted for most of the disagreement in the DTI and SMI metrics, and ABCD accounted for most of the disagreement in the DKI metrics (**Figure 4-5**).

Although inter-protocol variability exceeded test-retest variability (1-3%) for all metrics except RK (**Figure 1**), its magnitude was largely within the range reported for other quantitative MRI methods on a single-vendor platform. Reported average CVs range from 1.3% to 6.1% for T1, 3.4% to 4.1% for magnetization transfer ratio (MTR), and 3.4% to 5.9% for magnetization transfer saturation index (MTsat) (Karakuzu et al., 2022). An average CV of 2% has also been reported for T1 morphometry within the same scanner (Warrington et al., 2025). Together, these findings support the potential for pooling quantitative dMRI data across commonly used acquisition protocols, with b-value alignment improving agreement for several metrics.

### 4.1 Contributions of other acquisition parameters

Using our subject-matched data across various protocols, we quantified the impact of individual scan parameters on diffusion metrics. Within this study, we focused on TE, maximum *b*-value, and SNR, because (a) they accounted for most of the protocol variance, and (b) their effects could be estimated without inflation from collinearity in our data. Maximum *b*-value was the acquisition parameter most strongly associated with inter-protocol variability in the DTI diffusivities and the kurtosis metrics (**Figure 6**). However, contributions of SNR and TE cannot be ignored. SNR scales with voxel volume, but larger voxels increase partial-volume effects that may reduce the accuracy of diffusion metrics. TE can also alter the relative signal contributions of different microstructural compartments (Gong et al., 2020; Lampinen et al., 2019; Tax et al., 2021; Veraart et al., 2018). In this dataset, SNR had the second largest effects on the kurtosis metrics, whereas TE showed little association with any metric (**Figure 6**). No single acquisition parameter dominated the SMI metrics. TE, maximum *b*-value, and SNR accounted for most of the protocol variance in MK and AK but left a large proportion of the variance in *p*_2_ unexplained (**Table 2**).

Differences in diffusion gradient timing may also contribute to inter-protocol variability. Based on the available sequence information, the estimated gradient separations (Δ) ranged from approximately 34 to 46 ms, while gradient durations (δ) ranged from approximately 14 to 29 ms. Previous studies have demonstrated measurable dependence of diffusivity on both Δ and δ in human white matter, although these effects are relatively modest over the diffusion timing ranges typically used in clinical acquisitions (Fieremans et al., 2016; H.-H. Lee et al., 2018). We therefore expect differences in gradient timing to contribute relatively little to inter-protocol variability in conventional diffusivity metrics, although their effects on higher-order diffusion metrics remain uncertain. Further studies with independently varied acquisition parameters, including diffusion gradient timing, TR and acceleration factors, would be needed to isolate their individual contributions.

### 4.2 Maximum b-range and the effects of matching b-values

Protocols with higher maximum *b*-values showed greater disagreement with the other protocols. HCP-DA1 (*b*=0, 1500, and 3000 s/mm²) contributed most to the disagreement in DTI diffusivities, while ABCD (maximum *b-*value 3000 s/mm²) contributed most to the disagreement in kurtosis metrics (**Figure 4**). The compounding and decomposition analyses supported this pattern, as variability increased when these protocols were added and was most strongly associated with maximum *b*-value (**Figure 5**). DTI diffusivities were lower in protocols with a higher maximum single-shell *b*-value (**Figure 6**). Kurtosis metrics were lower in protocols with a higher maximum multi-shell *b*-value used for DKI and SMI estimation, but MK and AK also varied with the maximum DTI *b*-value (**Figure 6**). These findings suggest that the *b*-values of both the DTI and higher shells contributed to inter-protocol variability.

Matching *b*-values provided a direct test of this relationship and produced the largest reductions in variability for DTI diffusivities. Although previous work has shown that *b*-value selection can affect FA estimates (Kornaropoulos et al., 2022), FA showed little association with maximum *b*-value in this study and changed little after matching (**Figure 1**; **Table 3**). Matching also reduced variability in MK, AK, and *f*, but not in RK or *p*_2_, where it increased in most regions (**Figure 1**; **Table 3**). The high test-retest variability of RK made protocol-related changes difficult to distinguish from measurement noise, while *p*_2_ already had low inter-protocol variability (**Figure 1**).

The loss of data associated with the *b*-value matching analysis may also explain why some metrics did not improve. Advanced dMRI methods such as DKI and SMI require multiple *b*-value shells and sufficient diffusion directions to resolve fiber orientation and sensitivity to tissue microstructure (Jespersen et al., 2010; Novikov et al., 2019). Matching *b-*values removed about two thirds of the diffusion-weighted volumes from ABCD and few or none from the other protocols. Restricting the available *b*-values may reduce the precision of RK and *p*_2_ enough to offset gains in agreement.

Hence, for optimal precision, prospective studies would benefit from matching *b*-values to the same as the protocols they aim to combine with.

### 4.3 Regional heterogeneity

Protocol effects varied across WM regions, with the fornix and several projection tracts showing greater inter-protocol variability than regions such as the corona radiata (**Figure 2**). The regions with the greatest dispersion overlapped only partly with the ICC and Kendall’s *W* outliers (**Figure 3)**. This limited overlap reflects the different properties measured by absolute dispersion, agreement relative to inter-subject variance, and rank preservation. The high variability in the fornix is consistent with its small cross-sectional area and susceptibility to partial-volume contamination. However, *CV_protocol_* was only weakly related to region size, and smaller regions showed higher variability only for AK and *p*_2_ (**Supplementary figure S3**). Region size therefore does not fully explain the observed pattern. Regional differences may instead reflect interactions among local anatomy, measurement properties, noise, and acquisition parameters, consistent with previous reports of spatial variation in the sensitivity of anisotropy measures (Basser & Pierpaoli, 1996). Excluding the most variable regions produced little change in the overall findings, indicating that these regions did not drive the study-level results.

### 4.4 Implications for pooling multi-protocol data

Although no standard dMRI acquisition protocol exists, the field is increasingly emphasizing harmonization and compatibility (Bayer et al., 2022; Bento et al., 2022; Pinto et al., 2020; Stamoulou et al., 2022). Many studies adopt protocols modeled after large consortia, such as UK Biobank or ABCD, to facilitate comparability (Dean et al., 2024; Duff et al., 2022; Griffanti et al., 2022). The present findings provide quantitative guidance on which protocol features and combinations yield greater downstream compatibility under controlled conditions. For DTI metrics, protocols that share the same maximum *b*-value can be pooled with the least added variability, and restricting estimation to matched *b*-values reduced *CV_protocol_* by about half for RD and about two thirds for MD and AD (**Table 3**). Adding a multi-shell protocol whose maximum *b*-value differs from the others, here ABCD or HCP-DA1, introduced the largest increases in variability, and the compounding order identifies which cohorts can be combined first with the least added variability (**Figure 5**).

Most of the differences between protocols consisted of a constant offset across subjects, representing the type of systematic difference that post-hoc harmonization approaches such as ComBat are designed to correct. In pooled studies, however, each protocol is often associated with a different cohort, and these cohorts may differ systematically in characteristics such as age or clinical status. Consequently, protocol and cohort effects can become confounded, such that a harmonization method estimating protocol effects from the pooled data may also attribute cohort differences that coincide with protocol to the protocol effect. Our dataset here provides a benchmark of the same set of both young and old participants sampled across all protocols, making it ideal for development and testing of harmonization methods, and complementary to prior efforts that focused on test-retest (Lucignani et al., 2021) and inter-scanner variability only (Warrington et al., 2023, 2025).

### 4.5 Limitations

There are a few limitations that should be considered. First, data were acquired on a single scanner and vendor platform to isolate acquisition protocol variability independent of hardware and vendor effects. By holding these factors constant while systematically varying acquisition parameters, our design enables direct quantification of protocol driven variability and complements existing traveling heads resources focused on scanner and site effects. However, the use of a single scanner may limit generalization across vendors and introduce discrepancies relative to large-scale cohort data. Second, although this dataset includes a broad range of commonly used multi-shell protocols, it is not exhaustive of all acquisition schemes employed in large-scale neuroimaging studies. Even so, the coverage and distribution of acquisition parameters such as spatial resolution, *b*-value, and TE are representative of most multi-shell protocols. Third, the cohort consisted of healthy adults, so the findings may not generalize to patient cohorts or to the infants and young children included in lifespan studies. dMRI protocols for infants may include specific accommodations, and different *b*-value strategies have been evaluated for this age group, leading to protocols with their own range of *b*-values and sampling schemes (Howell et al., 2019). Water mobility in the neonatal and infant brain differs from that in adults, likely because of differences in tissue microstructure and diffusion barriers (Dudink et al., 2008; Jones et al., 2003; Mulkern et al., 2001; Tournier et al., 2020). The *b*-value effects estimated here in adults, and the benefit of matching *b*-values, may therefore not carry over to these populations. Finally, all protocols were acquired in a fixed order; consequently, protocol specific effects may be partially confounded by time dependent factors, such as participant fatigue, motion, or scanner drift.

Several aspects of the analysis should also be noted. *CV_ws_* was measured for one protocol (MESO-96), from two runs within a single session, and may differ for other protocols or across sessions (Lucignani et al., 2021). In addition, the tests of H1 and H2 used the 27 regions as paired observations, which are not fully independent within subjects; we therefore report the rank-biserial effect size alongside each *p*-value. Finally, we considered results may depend on processing choices, which we examined in supplementary analyses. Test-retest reproducibility improved with the advanced DESIGNER dMRI processing pipeline and parametric computation than with the HCP minimal processing pipeline (J. Chen et al., 2024; Glasser et al., 2013); **Supplementary Figure S1**). Registration was not the principal source of inter-protocol variability, as nonlinear deformations varied primarily between subjects rather than protocols, and *CV_protocol_* estimates remained largely consistent when recomputed using a shared subject-template registration (**Supplementary Figures S8-S9**).

### 4.6 Future Work

Future work can expand on these findings and address these limitations by incorporating publicly available datasets spanning different vendors and scanner models (Warrington et al., 2023, 2025). This would enable evaluation of combined protocol and hardware variability. Applying post hoc harmonization approaches to the present dataset would clarify how methods such as ComBat (Fortin et al., 2017; Hu et al., 2023; Johnson et al., 2007) or RISH (Mirzaalian et al., 2015) affect inter-protocol variability and biological rank preservation when scanner and site effects are already controlled. Previous work has found that ComBat improves inter-scanner variability, but inconsistencies in biological rank remained (Warrington et al., 2023). Because the protocol differences found here were largely shifts in level with preserved ranking, this dataset provides a direct test of whether harmonization reduces *CV_protocol_* without altering rank agreement. At the acquisition stage, future protocol designs and multi-cohort studies could target *b*-value alignment, for example by including a common 1000 s/mm² shell for DTI estimation, since matching *b*-values reduced inter-protocol variability for the metrics most sensitive to *b*-value. Extending the population to younger volunteers may help establish whether the same acquisition parameters account for inter-protocol variability across the lifespan. Additionally, alternative analytic approaches can be explored to determine whether our findings generalize across other methodological choices. These include tract-based analyses such as bundle analytics (Chandio et al., 2020) and computing DTI through kurtosis estimation (Veraart et al., 2011). Including the number of diffusion directions and voxel size in the decomposition, and estimating test-retest variability across sessions and protocols, would address the remaining gaps identified above.

## 5. Conclusion

This dataset allowed us to examine compatibility between large-scale, multi-shell dMRI protocols independent of biological, scanner, and site differences. This study demonstrates modest variability and largely preserved subject rankings across commonly used multi-shell dMRI acquisition protocols under controlled single-scanner conditions. FA showed particularly high stability, while aligning *b*-values substantially improved agreement for DTI diffusivities and reduced variability in MK, AK, and f. Although protocol related differences generally exceed test-retest variability, these findings provide a quantitative foundation for combining dMRI data across protocols and identify *b*-value alignment as a practical strategy for improving comparability. Together, these findings may inform efforts to select protocols, harmonize measurements, and construct normative models across large-scale neuroimaging datasets.

## Data and Code Availability

Data and code available upon request.

## Declaration of the Use of AI

Artificial intelligence tools were used to assist with language editing and to support code development. All analyses, methodological decisions, and interpretations were conducted and verified by the authors.

## Author Contributions

EF, JV and DSN conceived the study. JWJ wrote the manuscript, ran the analysis and visualizations for initial submission and revisions. JWJ, JC, and BAA were involved in image processing. VS, SM, NJ contributed to recruitment and data collection. EF, DSN, JV, SC, and RCL contributed technical expertise on image processing and quantification. EF and JV supervised the work and revisions. All authors contributed to the experimental design, the interpretation of the results, and editing of the final manuscript. All authors accept the final responsibility to submit for publication and take responsibility for the contents of the manuscript.

## Funding

This work has been supported by NIH NINDS (R01 NS088040), NIBIB (R01 EB027075, K99 EB03608), NIA (R33AG087904,RF1AG098531), the Irma T. Hirschl foundation, and was performed under the rubric of the Center for Advanced Imaging Innovation and Research (CAI2R, www.cai2r.net), an NIBIB National Center for Biomedical Imaging and Bioengineering (P41 EB017183).

## Competing Interests

J.V., D.S.N. and E.F. are co-inventors of patent US20180120404A1 (licensed) describing the denoising technology, and patent US20160343129A1 describing SM parameter estimation, that have been used in this study. D.S.N., E.F., and B.A.-A. are shareholders at Microstructure Imaging, Inc. that has licensed the patent.

## APPENDIX

**Appendix Figure A1.**
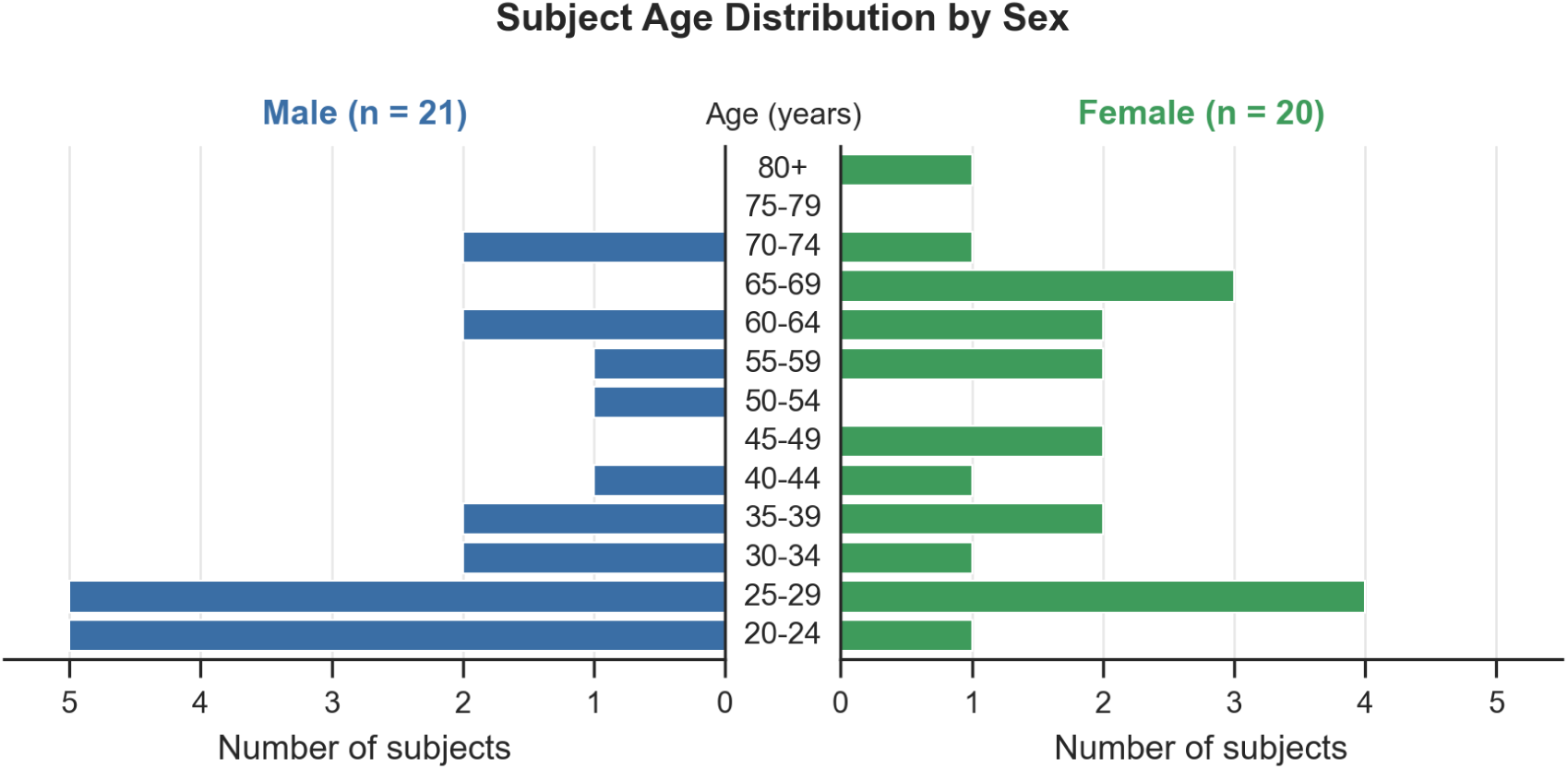
Age distribution of study participants stratified by sex. Horizontal bars represent the number of subjects (n=41) within 5 year age bins, with males in blue and females in green.

**Appendix Figure A2.**
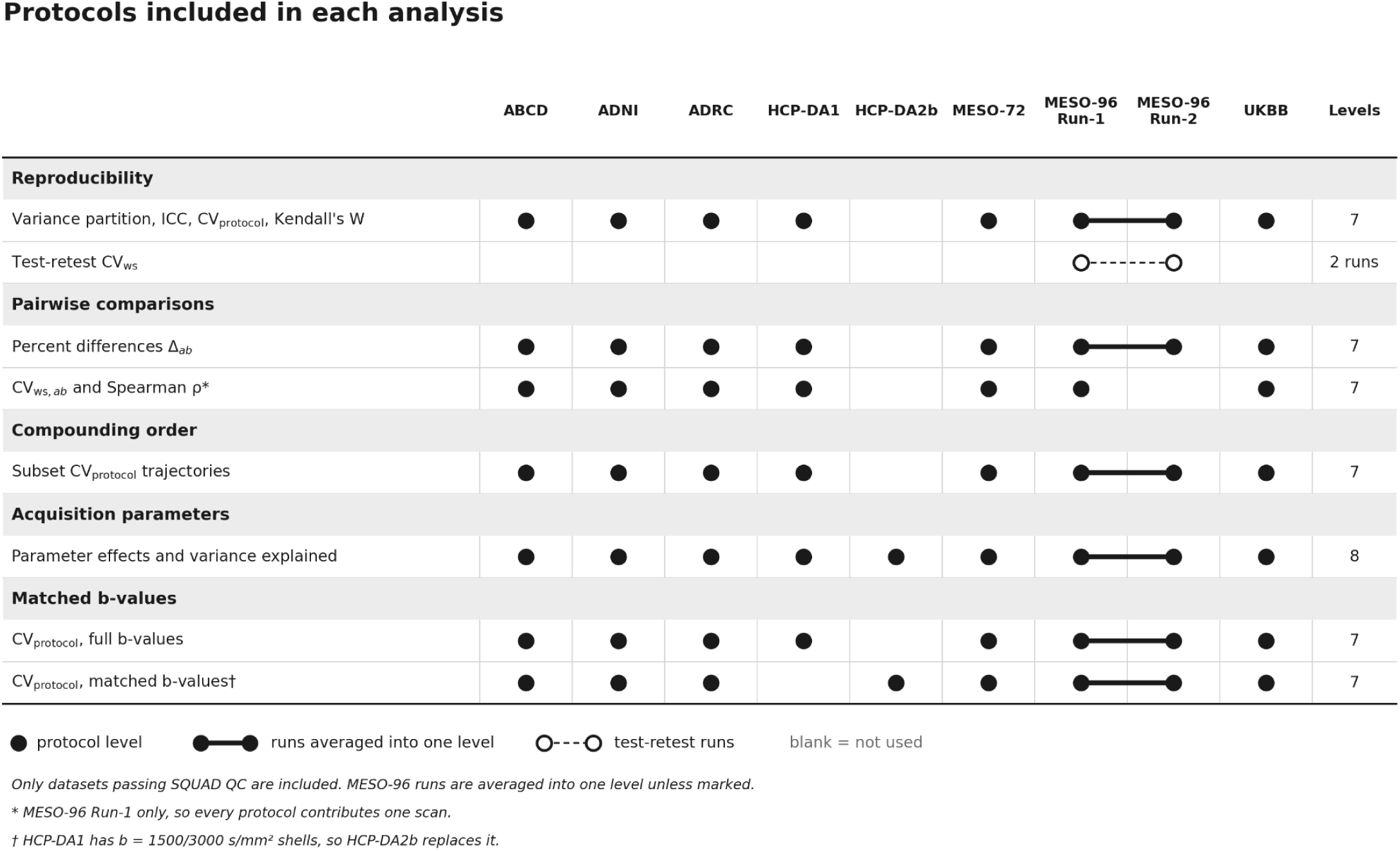
*Protocols included in each analysis*. Rows are the analyses described in Sections 2.6.2 to 2.6.7, grouped by section. Columns are the nine acquisitions. A filled circle marks a protocol entered as its own level. Filled circles joined by a solid bar mark MESO-96 Run-1 and Run-2 averaged into a single level. Open circles joined by a dashed line mark the two MESO-96 runs used as test-retest replicates for *CV_ws_*. Analyses marked * use MESO-96 Run-1 only, so that every protocol contributes one scan. Levels is the number of protocol levels in each analysis. HCP-DA2b enters the acquisition-parameter analysis as an eighth protocol. It also replaces HCP-DA1 in the matched *b*-value comparison (†), because HCP-DA1 was acquired with *b* = 1500 and 3000 s/mm² shells. Only datasets that passed SQUAD QC are included.

**Appendix Figure A3.**
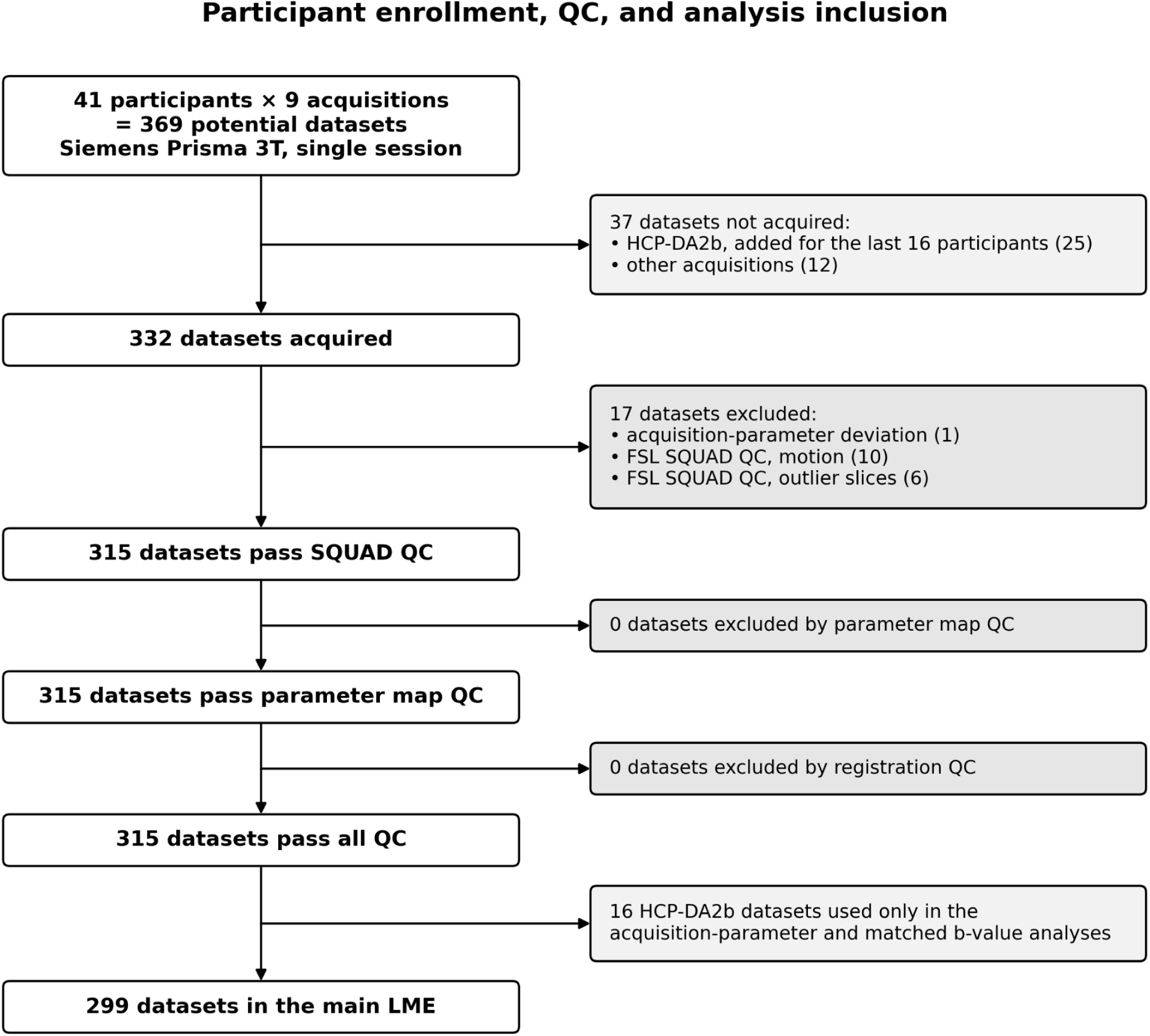
*Participant enrollment, quality control, and inclusion in the main LME*. Forty-one participants and nine acquisitions gave 369 potential datasets, of which 332 were acquired. FSL SQUAD group-level QC excluded datasets whose motion or outlier-slice percentage exceeded the group mean by more than 3 SD; datasets flagged on both criteria are counted under motion. Visual QC of parameter maps and registrations excluded no further datasets. The 16 HCP-DA2b datasets enter only the acquisition parameter and matched *b*-value analyses. The remaining 299 datasets gave 131,859 ROI × metric cells (49 JHU regions × 9 metrics). Of these, 51 (0.04%) had no valid value, 45 of them because the fornix contained no voxels in native space. Each missing cell is absent only from the corresponding region × metric estimation. Main LME in this case, includes the 7 main protocols (excluding HCP-DA2b).

**Appendix Table A1.**
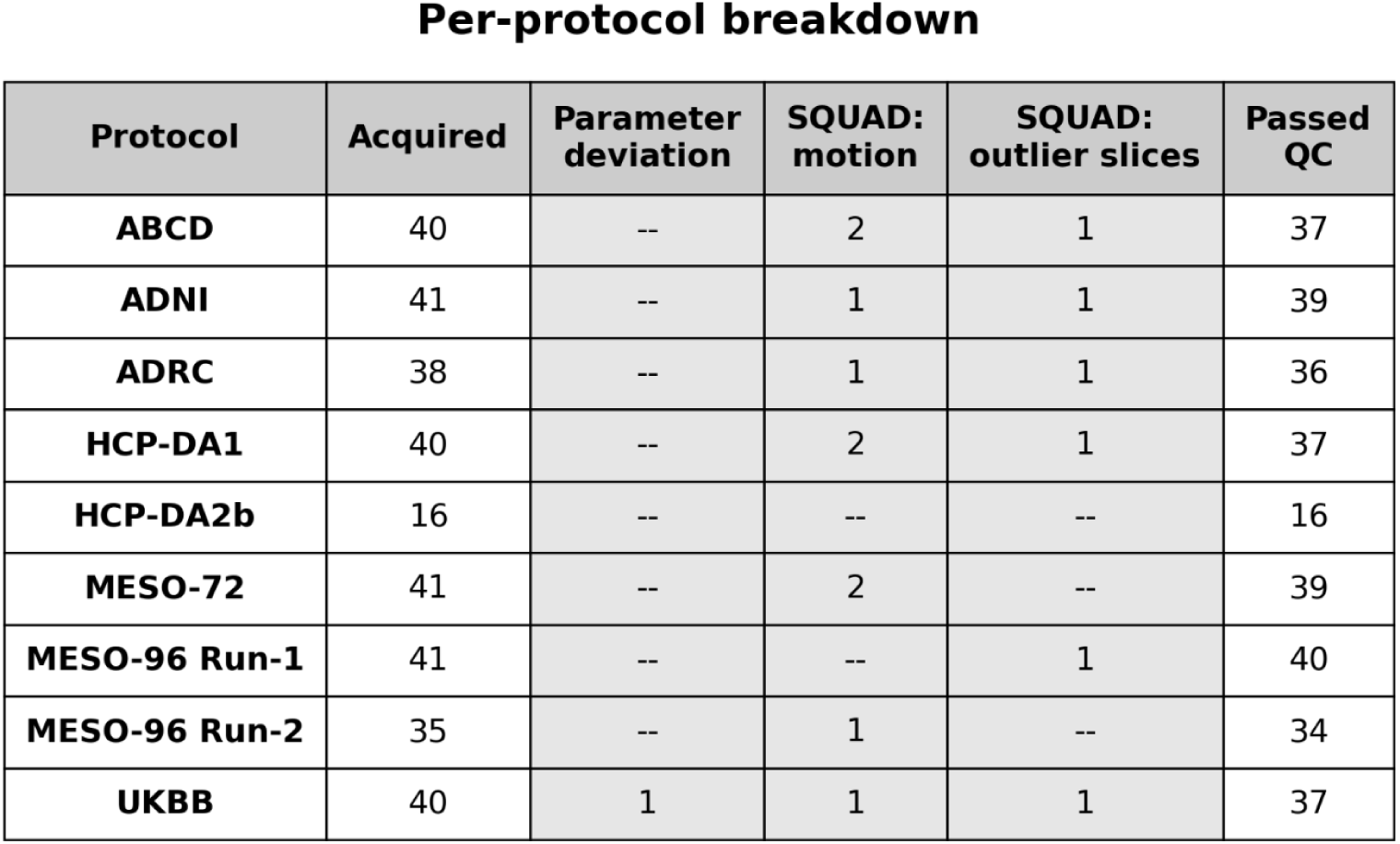
*Dataset counts by protocol*. Acquired is the number of datasets collected. SQUAD: motion and SQUAD: outlier slices are the datasets excluded by FSL SQUAD QC for each reason. Passed QC is the number of datasets retained. ROI × metric cells is the number of retained datasets × 49 JHU regions × 9 metrics. Cells unavailable is the number of those cells with no valid value. Grey columns count datasets or cells removed, and white columns count those retained. MESO-96 runs are listed separately but are averaged into one level in the LME. HCP-DA2b is used only in the acquisition-parameter and matched *b*-value analyses. “--” indicates zero.

