## Supplementary Material for "Inter-protocol reproducibility of multi-shell diffusion MRI metrics across major neuroimaging study protocols"

### *Image Processing*

MESO-96 test-retests runs were processed with both DESIGNER's advanced processing pipeline and Human Connectome Project's minimal diffusion processing pipeline (Glasser et al., 2013) which includes *b*=0 normalization, subject motion and geometric distortions from susceptibility and eddy current effects. Test-retest reproducibility improved with the advanced DESIGNER dMRI processing pipeline and parametric computation **(Supplementary Figure S1).**

### *Statistical Analyses*

All statistical analyses were performed in R v4.5.2 using linear mixed effects models (LME) (Laird and Ware, 1982) fit by restricted maximum likelihood (REML) with the *lme4* package (v2.0-1), with degrees of freedom and p-values for fixed effects obtained bySatterthwaite's approximation as implemented in lmerTest (v3.2-1). Estimated marginal means and protocol contrasts were computed with emmeans (v2.0.3), marginal and conditional coefficients of determination with MuMIn (v1.48), and confidence intervals for variance-derived quantities by parametric bootstrap using lme4::bootMer. Rank-based concordance statistics were generated in Python v3.11.8 (NumPy/Pandas/SciPy/Matplotlib) from the exported model tables and from the same quality-controlled ROI table used for model fitting. The Wilcoxon signed-rank tests and rank correlations were performed in Python v3.11.8 (NumPy/Pandas; SciPy).

### *Data Preparation*

**Appendix Figure A3 traces** the 369 potential datasets (41 participants × 9 acquisitions) to the 315 that passed quality control, and Table A1 gives the counts by protocol. HCP-DA2b was added partway through the study and was acquired in the last 16 participants only. The 16 SQUAD exclusions came from 10 participants, one of whom contributed six. For the main LME the two MESO-96 runs were averaged within participant (33 participants had both runs and eight had one), so all 41 participants contributed to that protocol.

Three representations of the quality-controlled MESO TE = 96 data were constructed, each matched to the class of statistic it enters. The run average was used for mean-based quantities (model fitting, grand mean, $CV_{protocol}$, ICC, variance partitioning). Run 1 alone was used for spread-based, pairwise quantities ( Spearman's ρ and the within-subject inter-protocol CV), so that averaging across runs does not artificially inflate agreement between MESO TE = 96 and other protocols. Both runs together were used to estimate the test-retest floor.

### *Linear Mixed-Effects Model*

inter-protocol statistics reported in this work i.e. variance partitioning, ICC, grand means, pairwise contrasts, and acquisition-parameter attribution, derive from a single linear mixed-effects model fit separately for each metric by ROI combination by restricted maximum likelihood (REML). For the ROI metric value, $y_{ij}$, of subject, *i*, under protocol, *j*,

$$y_{ij} = \mu_j + \beta_1(Age_i - \overline{Age}) + u_i + \varepsilon_{ij} ,$$

where $\mu_j$ is the fixed mean of protocol *j* at the sample mean age, $\overline{Age}$, $\beta_1$ is the common age slope, $u_i \sim N(0, \sigma^2_{subject})$ is a subject-specific random intercept, and $\varepsilon_{ij} \sim N(0, \sigma^2_{residual})$ is the residual, with $u_i$ and $\varepsilon_{ij}$ mutually independent. Age enters as a covariate, so that protocols are compared at a common age and the fitted protocols means, $\hat{\mu}_j$, are age adjusted. Sex was not included because it is constant within each participant across protocols and therefore cannot confound the primary within-participant protocol contrasts. Because the protocols were deliberately selected acquisition designs rather than a random sample from a larger population, protocol was modeled as a fixed effect.

#### *Variance Decomposition and Inter-Protocol Reliability*

The subject and residual variances were estimated directly by the mixed model. Because protocol was modeled as a fixed effect, the inter-protocol variance was instead derived from the spread the fitted protocol means around the grand mean. The grand mean, $\overline{Y}$, is the unweighted average of *k* fitted protocol means (*k* = 7, or 8 where noted). Every CV reported in the manuscript uses a pooled grand mean as its denominator, $\overline{Y}_{pooled} = (\overline{Y}_{full} - \overline{Y}_{matched})/2$: the mean of the grand means from the full b-range and matched b-value models (see Matched b-value Comparison), computed for each metric and region. For analyses on hemisphere-averaged data, the left and right values were averaged. This shared denominator ensures CVs are expressed on the same percentage scale.

Each fitted mean carries ordinary estimation error that inflates this spread even when protocols do not differ, so each expected error contribution was subtracted and the result floored at zero,

$$\hat{\sigma}^2_{protocol} = max\left(0, s^2_{\mu} - \nu\right),$$

Where $s^2_{\mu}$ is the observed variance of the *k* protocol, means and $\nu$ is the mean estimation-error variance of those means, taken from the model's covariance of the fixed effects. The inter-protocol CV was then computed as $CV_{protocol} = 100 \times \hat{\sigma}_{protocol} / \overline{Y}_{pooled}$. To assess whether regional differences in $CV_{protocol}$ reflect region size, $CV_{protocol}$ was correlated with the mean number of voxels retained in each region after voxelwise outlier removal, across the 27 regions and separately for each metric.

Inter-protocol absolute agreement was quantified with the two-way, fixed-rater intraclass correlation coefficient for single measurements, ICC(A,1) (McGraw & Wong, 1996; Shrout & Fleiss, 1979),

$$ICC = \frac{\hat{\sigma}^2_{subject}}{\hat{\sigma}^2_{subject} + \hat{\sigma}^2_{protocol} + \hat{\sigma}^2_{residual}}.$$

Placing the protocol variance in the denominator counts any systematic offset between protocols as disagreement, so the coefficient reflects how closely a single scan reproduces a single subject's value when the protocol is changed.

Rank-order agreement across all $k$ protocols was assessed with Kendall's coefficient of concordance, $W$ (Kendall & Babington Smith, 1939). Subjects were ranked within each protocol, and $W$ was computed from the sum of each subject's ranks across protocols. It ranges from 0 (no agreement) to 1 (identical ranking). $W$ was computed from subjects with data under all protocols, using the MESO-96 run average. Unlike the ICC, $W$ is unaffected by systematic offsets between protocols.

Test-retest variability was estimated from the two MESO-96 acquisitions, and used as a proxy for baseline test-retest noise. For each participant with paired acquisitions, the within-subject standard deviation was calculated as $s_i = \frac{|x_{i,1} - x_{i,2}|}{\sqrt{2}}$, where $x_{i,1}$ and $x_{i,2}$ are the average ROI summary values from run 1 and run 2, respectively, and pooled across participants as the root-mean-square, $\hat{\sigma}_{ws} = \sqrt{\frac{1}{n}\sum_{i=1}^{n} s_i^2}$ , to compute the estimated standard pooled within-subject standard deviation for replicate measurements (Luque Laguna et al., 2020). Participants with both runs of the MESO-96 protocol where both scans passed QC were included (n=33). The within-subject coefficient of variation ($CV_{ws}$) was computed as, $CV_{ws} = 100 \times \hat{\sigma}_{ws} / \overline{Y}_{pooled}$ , where $\hat{\sigma}_{ws}$ was calculated directly from the paired scans without fitting the LME. Taking the difference between the two scans for each participant removes stable differences between participants, including differences associated with age. The resulting measure reflects variability that remains when the same participant is scanned twice. Age-related differences in the amount of measurement variability were not explicitly modeled.

The same within-subject estimator was applied to every pair of protocols. For protocols $a$ and $b$, and each subject scanned under both, the within-subject standard deviation was calculated as $s_i = \frac{|x_{i,a} - x_{i,b}|}{\sqrt{2}}$, where $x_{i,a}$ and $x_{i,b}$ are the ROI summary values for protocols $a$ and $b$, and pooled across participants as the root-mean-square $\hat{\sigma}_{ws,pair} = \sqrt{\frac{1}{n}\sum_{i=1}^{n} s_i^2}$ . The corresponding within-subject CV between protocols was computed as $CV_{ws,pair} = 100 \times \frac{\hat{\sigma}_{ws,pair}}{\hat{\mu}}$. Unlike the test-retest case, the two measurements come from different protocols, so this difference reflects both any systematic offset between the protocols and residual within-subject noise; it therefore measures total pairwise disagreement rather than noise alone. In $CV_{ws,pair}$ computations, only run 1 was used here, so that every protocol contributes a single scan with the same level of scan noise; averaging the two runs would lower MESO-96's noise and make it appear to agree more closely with the other protocols, whereas in the LME the run average is appropriate because averaging leaves the expected protocol mean unchanged.

#### *Pairwise Protocol Comparisons*

To determine which specific protocols drive the excess variability identified in $CV_{protocol}$ across all protocols, all pairwise contrasts were obtained from the same fixed-effects model. For every protocol pair ($a$, $b$), the age-adjusted mean difference from the LME model yields an estimated signed percent difference,

$$\Delta_{ab} = 100 \times \frac{(\hat{\mu}_a - \hat{\mu}_b)}{\overline{Y}_{pooled}},$$

which is antisymmetric by construction ($\Delta_{ab} = -\Delta_{ba}$). The percent difference between protocol pairs are determined from metric values reported from the WM mask (**Supplementary Material Figure 5**).

For each protocol pair, $CV_{ws,pair}$ and Spearman's rank correlation ($\rho$) were computed across the subjects scanned under both protocols. Spearman's $\rho$ was computed on unadjusted ROI values and measures whether two protocols rank subjects in the same order and is unaffected by systematic offsets between protocols. Where low $\rho$ indicates protocols do not preserve differences between subjects, high percent differences or $CV_{ws,pair}$ and high spearman's $\rho$ indicate systematic value differences between protocols.

#### *Protocol Compounding-Order Optimization*

Since $CV_{protocol}$ depends on which protocols are pooled, the change in the inter-protocols variability was tracked as protocols were added one at a time, pooled in an order that minimizes variance accumulation. To understand how inter-protocol variability accumulates as acquisition protocols are pooled into a combined dataset, we performed a compounding analysis. Starting from a pair of protocols, we sequentially add one protocol at a time and track how the $CV_{protocol}$ increases at each step. Orderings that keep this variability low across all steps identify protocol combinations that are most compatible for pooled analysis. For each subset ($S$) of the seven main protocols (ABCD, ADNI, ADRC, MESO-96, MESO-72, UKBB and HCP-DA1), $\hat{\sigma}_{protocol}(S)$ was computed using the subset's fitted means and their estimation-error term $\nu(S)$, floored to zero. The fitted means and their covariance came from the full seven-protocol LME fit for the given ROI and metric. $CV(S) = 100 \times \hat{\sigma}_{protocol}(S) \,/\, \overline{Y}_{pooled}$ . $\overline{Y}_{pooled}$ is fixed across all subsets, so changes between steps reflect only the change in spread between protocols

All 2,520 unique orderings (7! permutations, halved because the order of the first two protocols is interchangeable) were evaluated. For each ordering the trajectory $\{CV(S_2), \dots, CV(S_7)\}$ was computed, where $S_q$ contains the first $q$ protocols. Orderings were ranked by the Frobenius norm across the nine metrics and six compounding steps,

$$||CV||_F = \sqrt{\sum_{s=2}^{7} \sum_{m} CV_m(S_q)^2},$$

where $m$ indexes metric. The ordering with the smallest norm was considered optimal. Ranking by the mean of the per-step norms selected the same ordering in every analysis. This norm captures the total variability accumulated across all steps and all metrics in a single scalar, giving preference to orderings that keep variability low throughout the entire accumulation sequence rather than only at individual steps. The search was repeated within each metric family (DTI, DKI, SMI) and with the three families weighted equally.

The analysis was performed in total WM and repeated in 27 regions (hemispheres averaged. To verify that the WM findings were not an artifact of spatial averaging, we repeated the analysis at the regional level across JHU atlas regions. ROI based results were summarized as the mean $CV_{protocol}$ across regions at each compounding step, with uncertainty estimated via a 200-replicate subject level bootstrap (resampling subjects with replacement and refitting the model on each replicate). The 95% bootstrap confidence intervals of the mean are shown as shaded bands (**Supplementary Figure S6**).

To identify which metrics drove the overall increase in variability at each compounding step, we decomposed the aggregate variance into per-metric contributions. At each step, the total stacked height equals the sum of squared CV values across metrics, with each metric's share shown as a segment of the stacked bar.

***Decomposition of Inter-Protocol Variance by Acquisition Parameters***

This analysis tested whether inter-protocol differences can be attributed to three acquisition parameters: TE, maximum *b*-value, and WM SNR. Eight protocols, the seven main protocols plus HCP-DA2b were utilized in the model and ran for all 27, hemisphere averaged JHU regions. Each acquisition parameter was z-scored across the eight protocols, given $z_j^{TE}$, $z_j^{b,DTI}$, $z_j^{SNR}$ for protocol *j*. Here $z_j^{b,DTI}$ is the maximum *b*-value in the DTI shell (1000 or 1500 s/mm²)

$$y_{ij} = \beta_0 + \beta_1(Age_i - \overline{Age}) + \beta_{TE} z_j^{TE} + \beta_{b,DTI} z_j^{b,DTI} + \beta_{SNR} z_j^{SNR} + u_i + \varepsilon_{ij}\,.$$

For DKI and SMI metrics, $\beta_b z_j^b$ was added, where $z_j^b$ is the maximum *b*-value used for DKI and SMI estimation (2000 or 3000 s/mm²), to account for the full range *b*-range in multi-shell data. Each coefficient was standardized by dividing it by the standard deviation of the metric across all scans in the region. For example, a standardized effect of 0.5 for TE therefore means that a protocol whose TE is one protocol-level standard deviation higher measures the metric 0.5 standard deviations higher, with the other parameters held fixed. The sign gives the direction of the effect, and the magnitude allows comparison across parameters and metrics. Medians and interquartile ranges of the standardized effects across the 27 regions are reported by metric.

The share of inter-protocol variance explained by the parameters was estimated with a linear regression (ordinary least squares) of the eight centered protocol means on the same *z*-scored parameters, with one observation per protocol:

$$\hat{\mu}_j - \overline{Y} = \gamma_0 + \gamma_{TE} z_j^{TE} + \gamma_{b,DTI} z_j^{b,DTI} + \gamma_{SNR} z_j^{SNR} + e_j.$$

where $z_j^b$ is $z_j^{b,DTI}$ for DTI metrics and $z_j^{b,DKI}$ for DKI and SMI metrics, and $e_j$ is the protocol level residual. $R^2$ from this regression is the fraction of the spread in the protocol means accounted for by the three parameters, jointly. The protocol variance was split into an explained part, $R^2 \times \hat{\sigma}^2_{protocol}$ , and an unexplained part $(1 - R^2) \times \hat{\sigma}^2_{protocol}$ . Both are reported as percentages of $\hat{\sigma}^2_{protocol}$, together with adjusted $R^2$, which corrects for the number of parameters relative to the number of protocols. A large unexplained portion of the protocol variance alludes to protocols differing for reasons outside of the selected acquisition parameters included in the model or effects that are not linear in these parameters.

#### *Matched b-value Comparison*

To test directly how maximum *b*-value contributes to inter-protocol variability, $CV_{protocol}$ was compared between two versions of computed parametric maps: full *b*-range and matched *b*-values. The full version used each protocol's complete acquisition (DTI: $b \leq 1500$ s/mm²; DKI and SMI: all acquired shells) for the seven main protocols. The matched version used $b = 0$ and 1000 s/mm² for DTI and $b = 0$, 1000, and 2000 s/mm² for DKI and SMI. In the matched version, HCP-DA1 was replaced by HCP-DA2b. For each version, the LME was refit and $\hat{\sigma}_{protocol}$ was computed. For each metric and region, $CV_{protocol}$ in both versions was expressed relative to the pooled grand mean, $\overline{Y}_{pooled} = (\overline{Y}_{full} + \overline{Y}_{matched}) / 2$, so that differences between versions reflect only differences in inter-protocol spread. If maximum *b*-value contributes to inter-protocol variability, $CV_{protocol}$ should be lower with matched *b*-values.

$CV_{protocol}$ from the two versions was plotted for each region and metric (hemispheres averaged, whole-WM mask excluded) against the identity line (**Supplementary Figure S7**). Points below the line indicate lower inter-protocol variability with matched *b*-values. A point was labeled as an outlier when either value exceeded the upper Tukey fence of the pooled values $(Q_3 + 1.5 \times IQR)$, or when the difference between versions fell outside $Q_1 - 3 \times IQR$ or $Q_3 + 3 \times IQR$ of all differences.

#### *Statistical Inference*

Two hypotheses were tested formally for each diffusion metric using the 27 JHU regions as paired observations, with hemispheres averaged and the whole-WM mask excluded. **H1:** inter-protocol variability exceeds test-retest variability, $\widehat{CV}_{protocol} > \widehat{CV}_{ws}$ . **H2:** matching *b*-values reduces inter-protocol variability, matched *b*-range $\widehat{CV}_{protocol}$ < full *b*-range $\widehat{CV}_{protocol}$ . Each hypothesis was tested using a one-sided Wilcoxon signed-rank test on the corresponding regional differences, with the exact null distribution (n = 27, no ties). Effect size was quantified using the matched-pairs rank-biserial correlation ($r_{rb}$), which ranges from −1 to 1, with a magnitude of 1 indicating that all regions differed in the same direction. The nine metric-specific tests for each hypothesis were treated as separate families and Bonferroni corrected independently, with statistical significance defined as an adjusted *p*-value, $p_{corr} < 0.05$. All other quantities were reported descriptively.

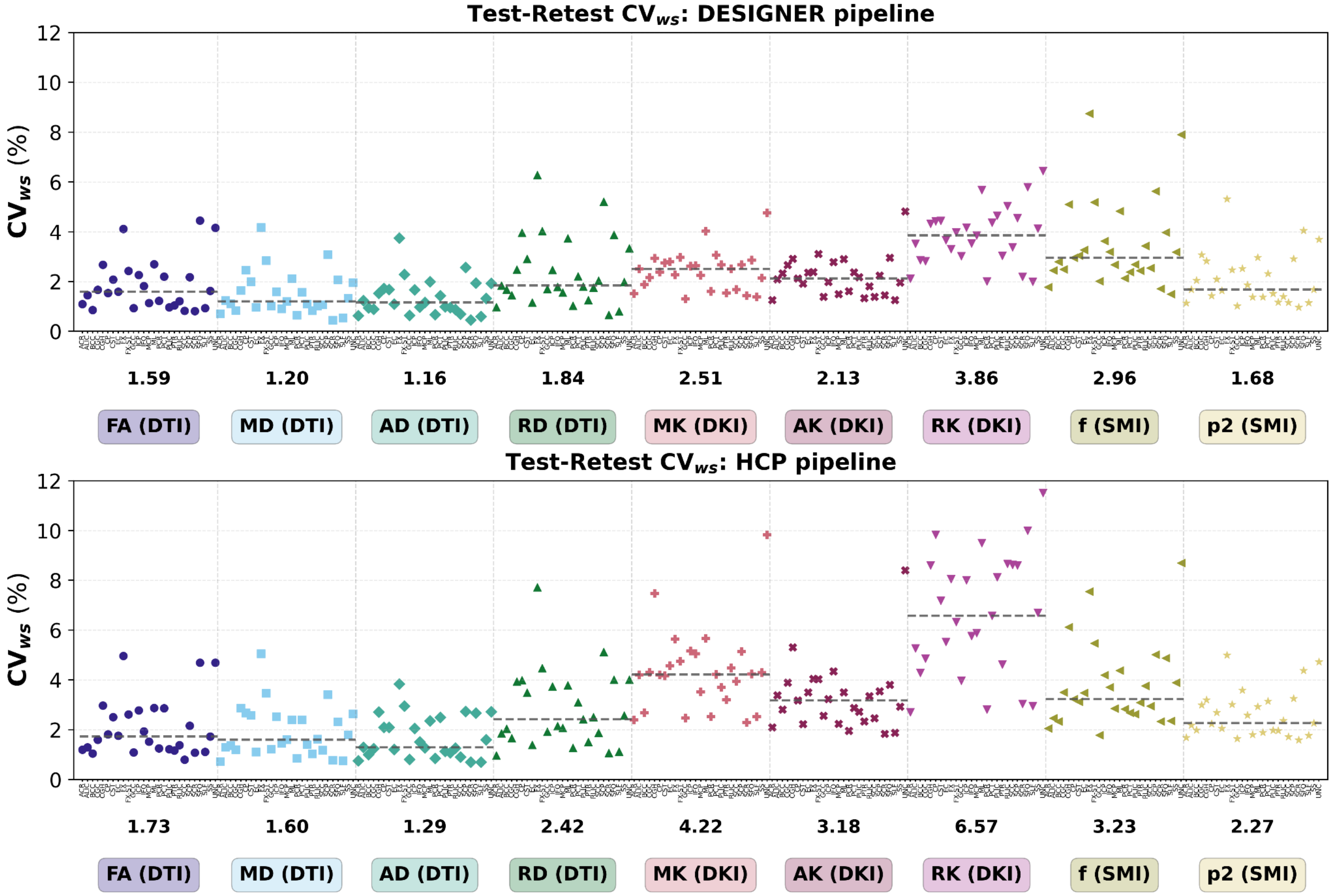


**Supplementary Figure S1**. *Test-retest repeatability of nine diffusion metrics under the DESIGNER and HCP processing pipelines.* Each point is the test-retest within-subject coefficient of variation ($CV_{ws}$) for one white matter region, from the two MESO-96 runs processed with DESIGNER (top) or the HCP pipeline (bottom). $CV_{ws}$ = 100 × σws / Ȳ, *where σ*ws = √(mean(d²)/2) and d is the run 1 − run 2 difference for each subject. Both pipelines use the same denominator: the intercept from a mixed-effects model fit to both pipelines and all protocols.. The ratio of the two pipelines' $CV_{ws}$ values is therefore the ratio of their within-subject SDs. Both pipelines use the same subjects in each region × metric (n = 31-33). Values for left and right regions are averaged across hemispheres. Dashed lines and the numbers below each group are the median $CV_{ws}$ across regions. Lower values mean better repeatability.

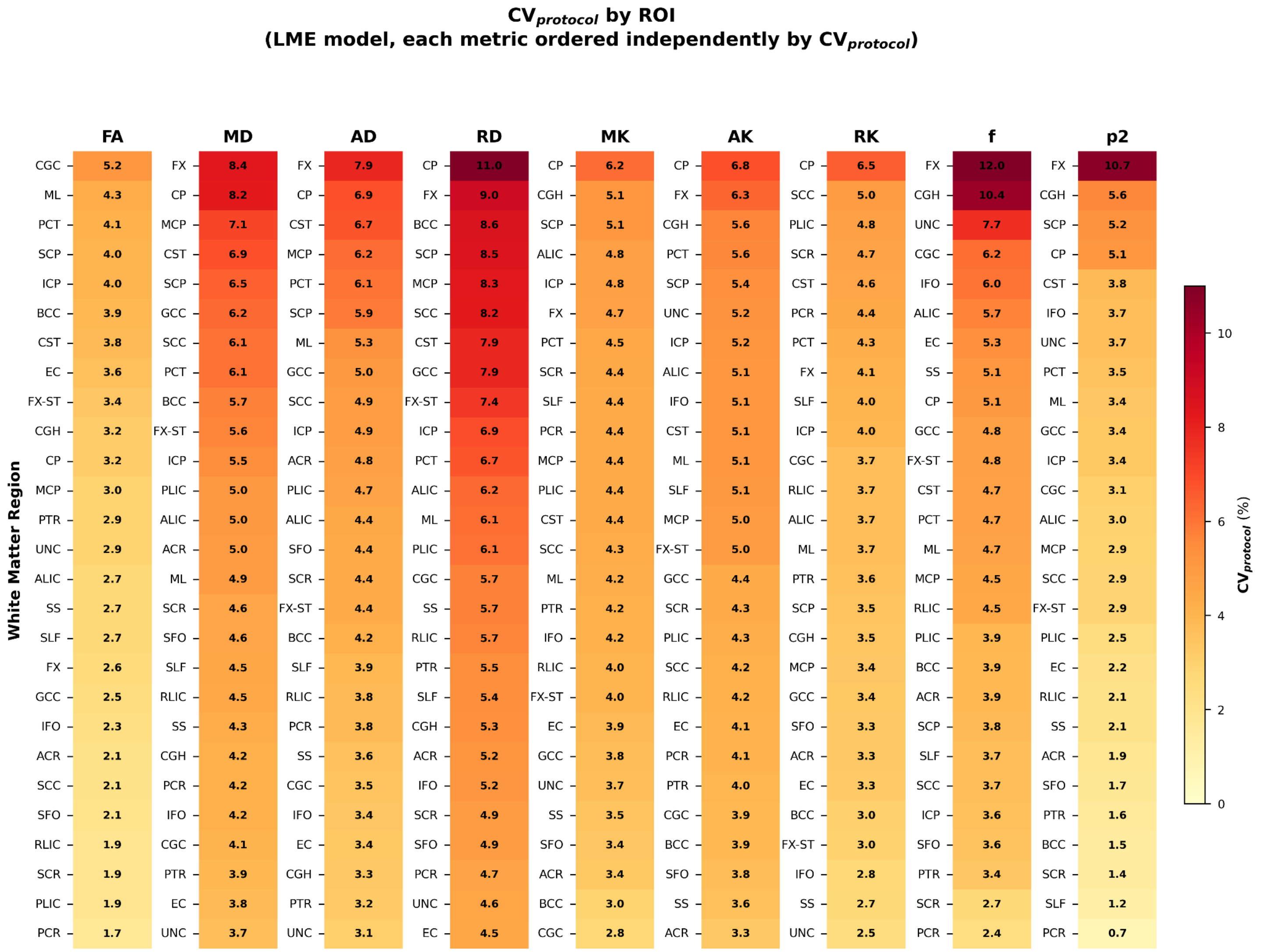


**Supplementary Figure S2.** *Regional inter-protocol coefficient of variation ($CV_{protocol}$) across metrics.* The $CV_{protocol}$ is reported for each JHU region (27 regions, averaged across hemispheres where applicable) within each metric. Regions are ranked from highest to lowest $CV_{protocol}$ within each diffusion metric, with light yellow being low $CV_{protocol}$ and dark red as a high $CV_{protocol}$. $CV_{protocol}$ was derived from LME results run for each JHU region and each metric.

**H1: $CV_{protocol}$ > test-retest $CV_{ws}$**

| Metric | $CV_{protocol}$ (%) | $CV_{ws}$ (%) | p (one-sided) | $p_{corr}$ | $r_{rb}$ |
|---|---|---|---|---|---|
| **FA** | 2.94 | 1.67 | < 0.001 | **0.005** | 0.69 |
| **MD** | 4.95 | 1.19 | < 0.001 | **< 0.001** | 1.00 |
| **AD** | 4.42 | 1.17 | < 0.001 | **< 0.001** | 1.00 |
| **RD** | 6.08 | 1.80 | < 0.001 | **< 0.001** | 1.00 |
| **MK** | 4.30 | 2.43 | < 0.001 | **< 0.001** | 0.98 |
| **AK** | 4.97 | 2.10 | < 0.001 | **< 0.001** | 1.00 |
| **RK** | 3.66 | 3.79 | 0.402 | 1.000 | 0.06 |
| **f** | 4.68 | 3.02 | < 0.001 | **< 0.001** | 0.83 |
| **p2** | 2.95 | 1.70 | < 0.001 | **0.002** | 0.73 |

*CVs are medians across 27 hemisphere-averaged JHU regions (whole-WM mask excluded). Exact one-sided Wilcoxon signed-rank test across regions; $p_{corr}$ = min(1, 9p), Bonferroni over the 9 metrics. Bold: significant ($p_{corr}$ < 0.05). Rank-biserial $r_{rb}$ > 0 supports H1.*

**Supplementary Table S1.** *One-sided tests of whether inter-protocol variability exceeds test-retest variability***.** For each diffusion metric, $CV_{protocol}$ and test-retest $CV_{ws}$ (MESO-96, run 1 vs run 2) were compared across 27 JHU regions (hemispheres averaged, whole white matter mask excluded) with an exact one-sided Wilcoxon signed-rank test. CV values are medians across regions. $p_{corr}$ is Bonferroni-corrected over the nine metrics, and bold marks $p_{corr}$ < 0.05. The $r_{rb}$ is the matched-pairs rank-biserial correlation, an effect size for how consistently regions differ in one direction. $r_{rb}$ > 0 indicates $CV_{protocol}$ > $CV_{ws}$, and $r_{rb}$ = 1 means every region differed in that direction.

**Variability across JHU WM regions vs whole WM**

| ***Across JHU ROIs (μ ± SD)*** | **FA** | **MD** | **AD** | **RD** | **MK** | **AK** | **RK** | **f** | **p2** |
|---|---|---|---|---|---|---|---|---|---|
| **ICC** | 0.66 (± 0.15) | 0.32 (± 0.17) | 0.33 (± 0.11) | 0.44 (± 0.17) | 0.36 (± 0.14) | 0.31 (± 0.11) | 0.59 (± 0.14) | 0.42 (± 0.17) | 0.55 (± 0.16) |
| **$CV_{protocol}$** | 3.00 (± 0.89) | 5.29 (± 1.29) | 4.67 (± 1.25) | 6.54 (± 1.65) | 4.22 (± 0.71) | 4.73 (± 0.84) | 3.79 (± 0.84) | 4.99 (± 2.11) | 3.16 (± 1.94) |
| **$CV_{ws}$** | 1.94 (± 1.10) | 1.48 (± 0.85) | 1.38 (± 0.74) | 2.28 (± 1.29) | 2.36 (± 0.76) | 2.16 (± 0.76) | 3.76 (± 1.12) | 3.53 (± 1.78) | 2.16 (± 1.11) |
| *Across JHU ROIs excluding FX, CP, CGH (μ ± SD)* | | | | | | | | | |
| **ICC** | 0.66 (± 0.14) | 0.32 (± 0.16) | 0.34 (± 0.11) | 0.44 (± 0.15) | 0.37 (± 0.12) | 0.32 (± 0.11) | 0.60 (± 0.13) | 0.44 (± 0.17) | 0.57 (± 0.16) |
| **$CV_{protocol}$** | 3.00 (± 0.94) | 5.08 (± 0.99) | 4.50 (± 0.99) | 6.30 (± 1.36) | 4.07 (± 0.57) | 4.54 (± 0.65) | 3.68 (± 0.68) | 4.47 (± 1.17) | 2.66 (± 1.05) |
| **$CV_{ws}$** | 1.82 (± 1.02) | 1.33 (± 0.68) | 1.25 (± 0.59) | 2.06 (± 1.09) | 2.35 (± 0.80) | 2.12 (± 0.79) | 3.74 (± 1.18) | 3.26 (± 1.47) | 2.01 (± 0.89) |
| *JHU white-matter mask* | | | | | | | | | |
| **ICC** | 0.64 | 0.24 | 0.19 | 0.34 | 0.32 | 0.27 | 0.55 | 0.50 | 0.54 |
| **$CV_{protocol}$** | 2.19 | 4.98 | 4.33 | 5.96 | 3.87 | 3.96 | 3.46 | 3.57 | 1.63 |
| **$CV_{ws}$** | 0.51 | 0.37 | 0.33 | 0.56 | 1.44 | 1.07 | 2.17 | 1.82 | 0.87 |

*LME model; 27 hemisphere-averaged JHU regions (24 excluding FX, CP, CGH); $CV_{ws}$ = MESO 96 run-1 vs run-2. Every CV uses that cell's LME grand mean.*

**Supplementary Table S2.** Agreement and variability in whole white matter and across JHU regions. Intra-class correlation coefficient (ICC), inter-protocol coefficient of variation ($CV_{protocol}$), and within-subject coefficient of variation ($CV_{ws}$) are reported for the whole JHU white matter mask and summarized across individual JHU regions as the mean (± SD). Regional summaries are provided both across all ROIs and across ROIs excluding three regions that show consistently high variability across metrics: the fornix (FX), cerebellar peduncles (CP), and the hippocampal portion of the cingulum (CGH).

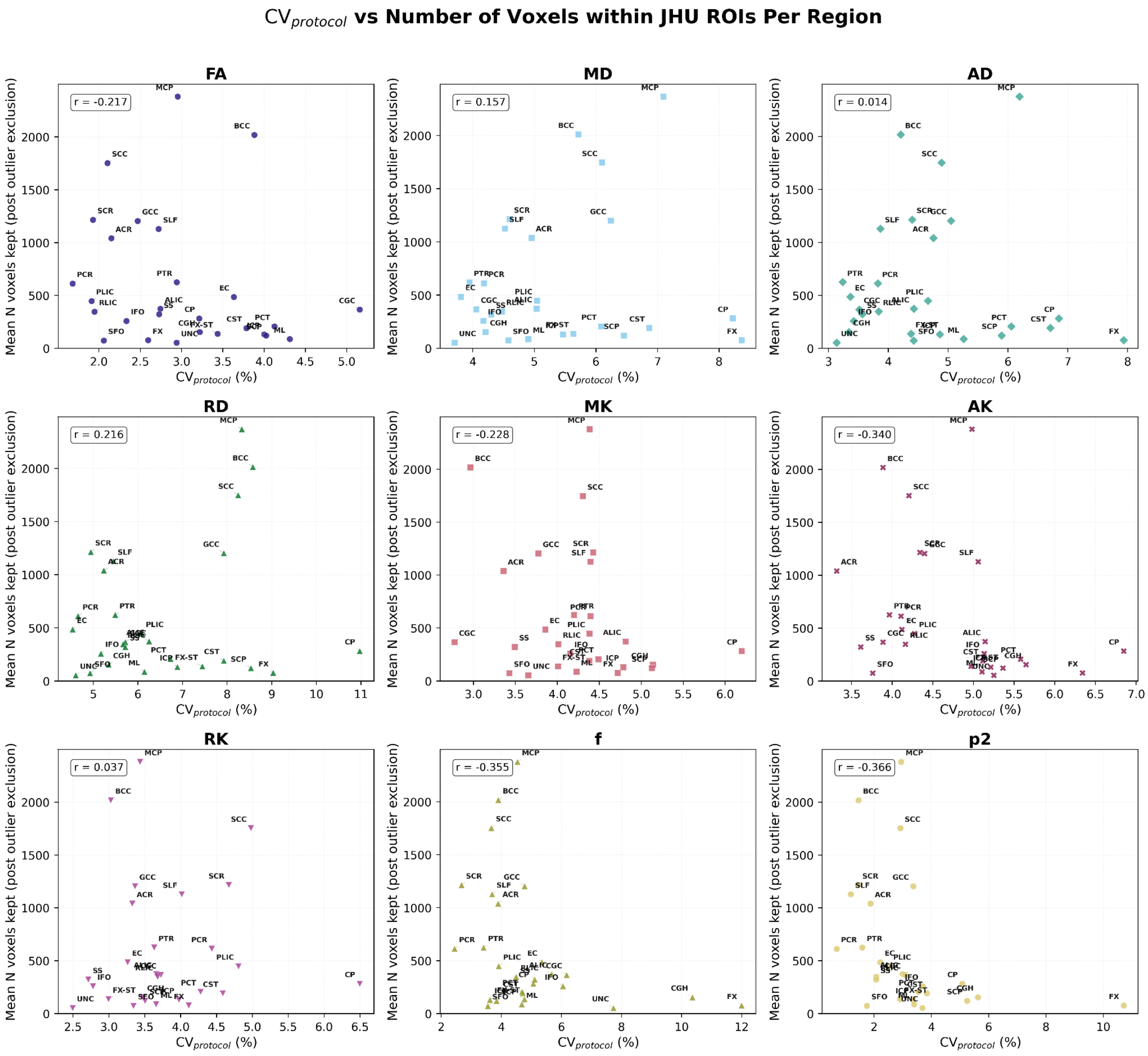


**Supplementary Figure S3.** Relationship between inter-protocol variability and ROI size across JHU white matter regions. Each panel displays one diffusion metric, with each point representing one hemisphere-averaged JHU white matter region. The x-axis shows the inter-protocol coefficient of variation (CV$_{protocol}$, %). The y-axis shows the average number of voxels retained per region after voxelwise outlier exclusion, averaged across subjects and acquisition protocols. Pearson correlation coefficients ($r$) are reported in each panel.

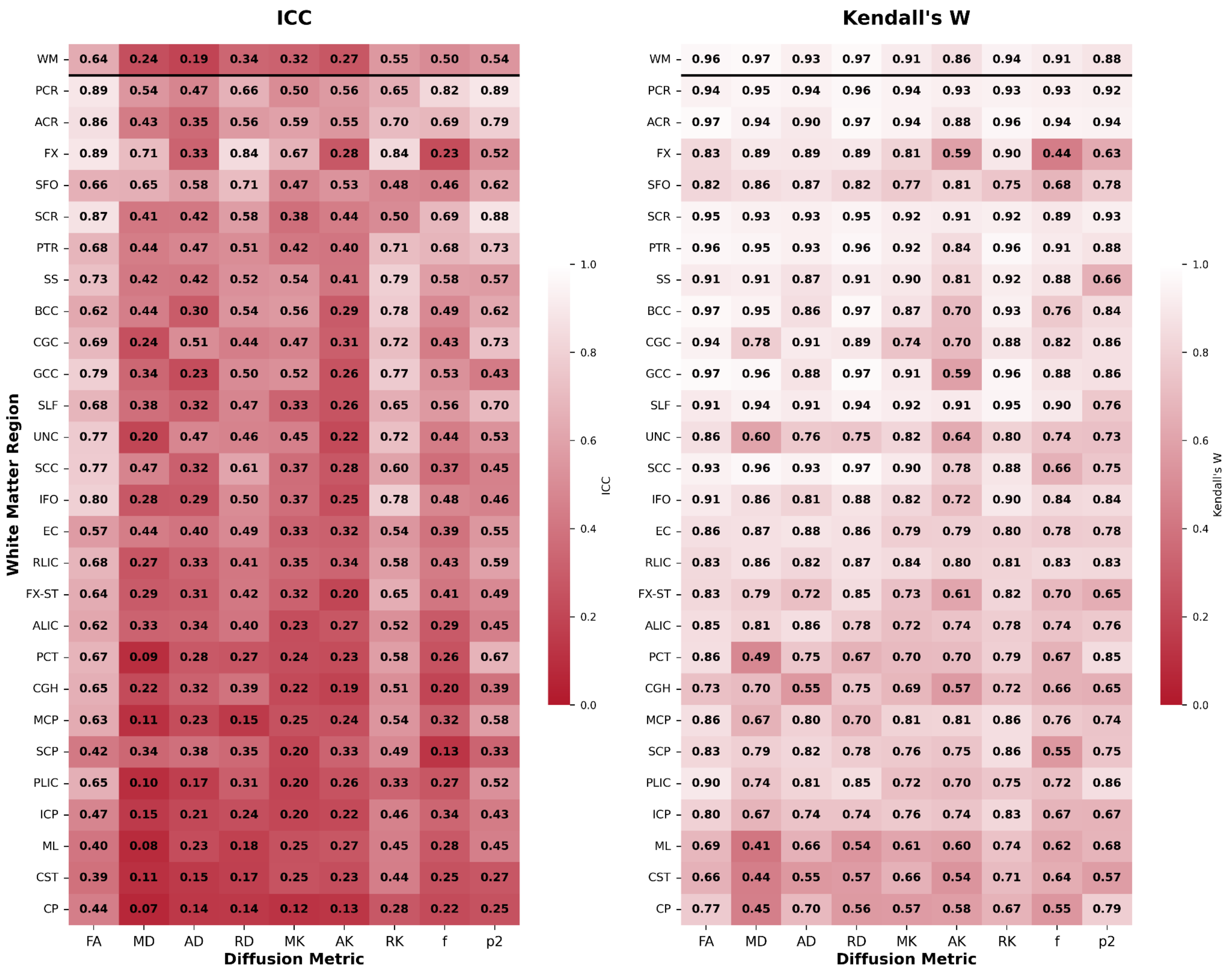


**Supplementary Figure S4.** *Cross-protocol reliability of diffusion metrics across JHU White Matter regions***.** Left: intraclass correlation coefficient (ICC, absolute agreement). Right: Kendall's W, which measures whether protocols rank subjects in the same order. Both panels use the same 0-1 color scale, from red (poor) to white (good). Rows are ordered by mean ICC across metrics. The whole white matter mask (WM) is shown above the black line. Values for left and right regions are averaged across hemispheres.

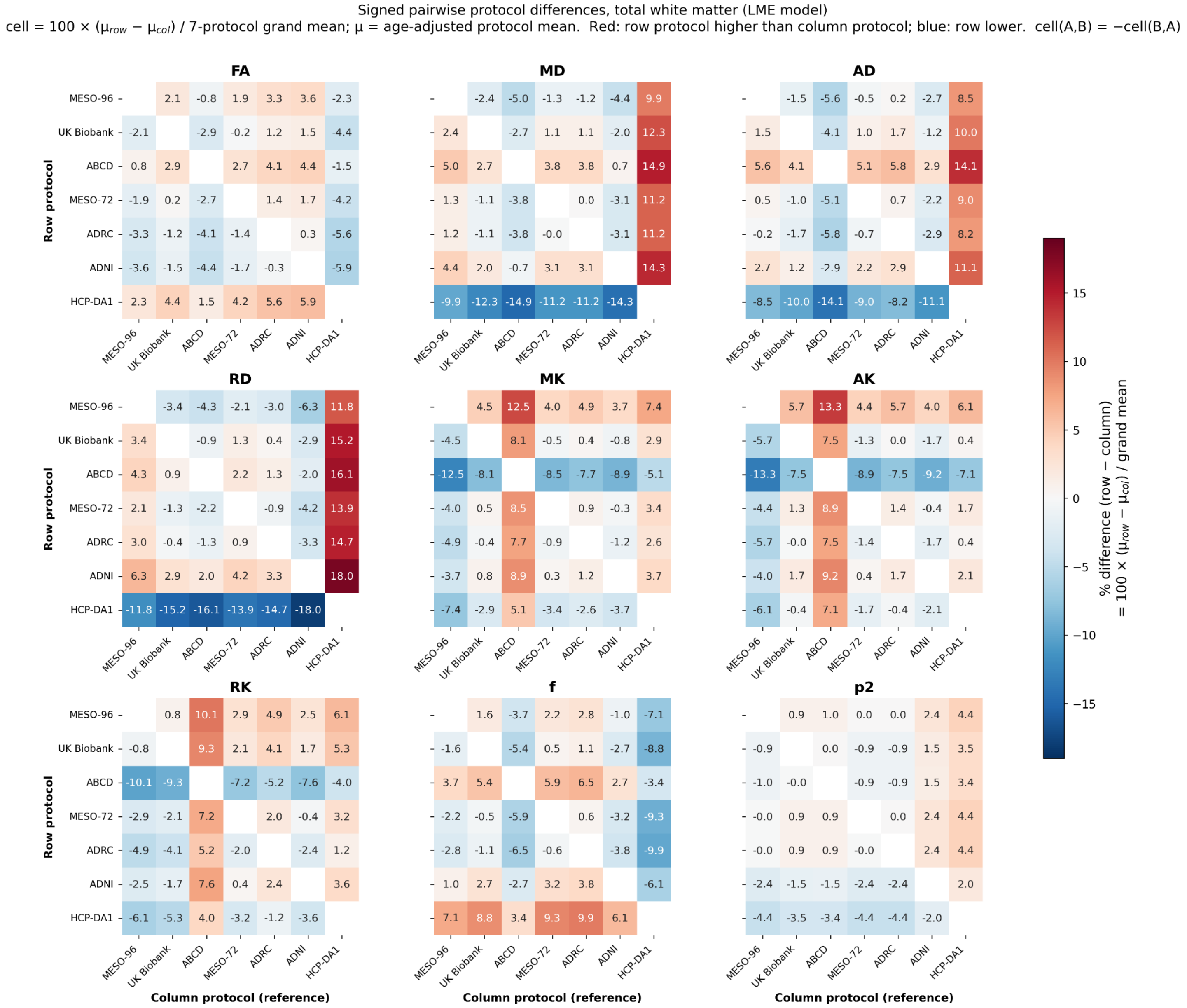


**Supplementary Figure S5**. *Signed differences in mean value between each pair of protocols in white matter.* Each panel is one diffusion metric, computed in the whole white matter mask. For each metric, a linear mixed-effects model was fitted with age as a fixed covariate, protocol as a fixed effect and a random intercept for each subject (41 subjects, 7 protocol levels). Each protocol's age-adjusted estimated marginal mean ( $\mu$) was then obtained. Each cell is the difference between the row and column protocols' marginal means, as a percentage of the pooled intercept: 100 × ($\mu_{row}$ − $\mu_{row}$) / $\overline{Y}$. The pooled intercept, $\overline{Y}$, is the unweighted average of the seven fitted protocol means. The two MESO-96 runs are averaged before model fitting. Red means the row protocol gives a higher value than the column protocol, and blue a lower one. The matrix is antisymmetric: cell (A, B) = −cell (B, A).

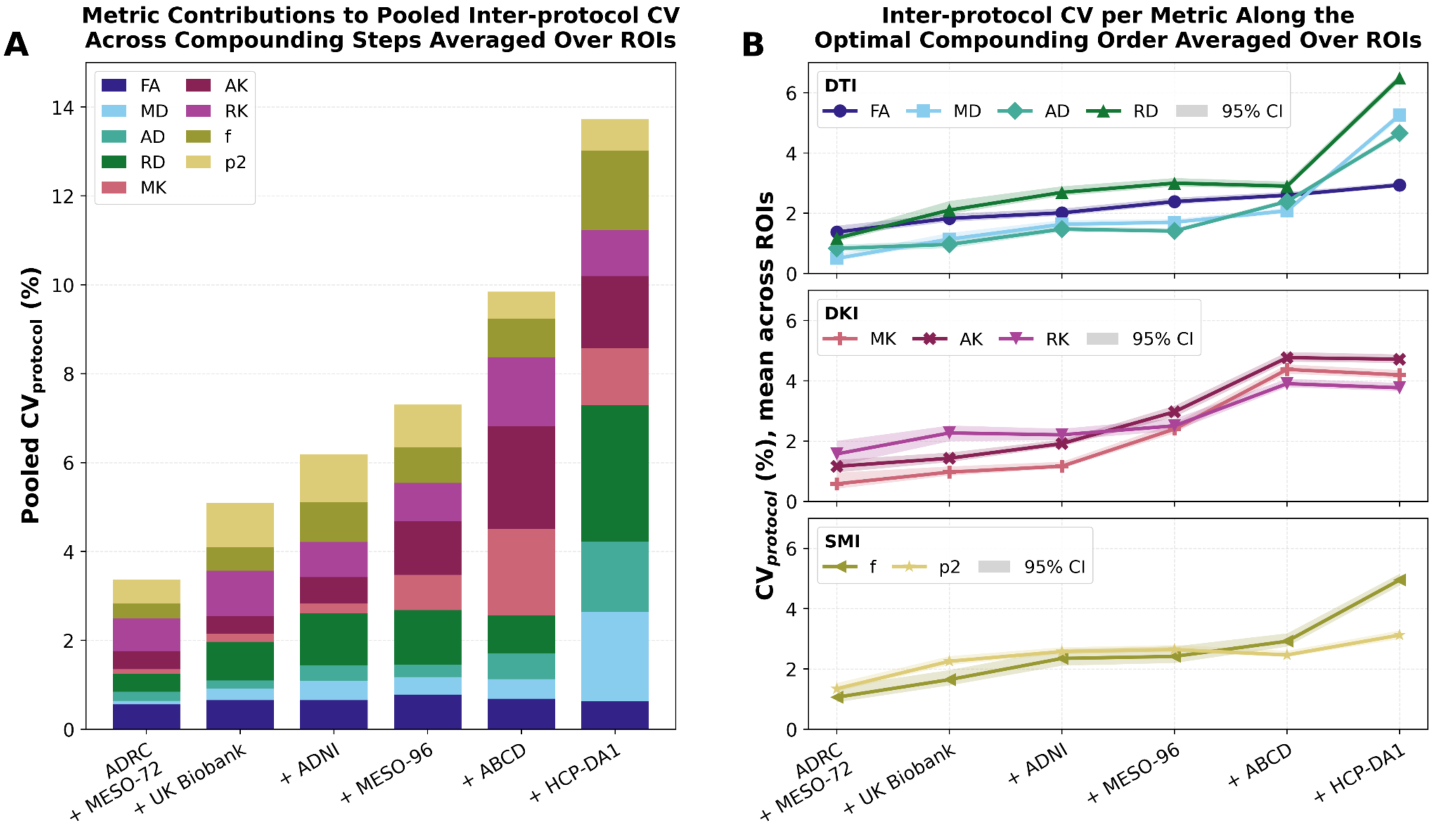


**Supplementary Figure S6**. *Accumulation of inter-protocol variability across individual white matter regions***.** $CV_{protocol}$ was computed in each of the 27 hemisphere-averaged JHU regions (whole-WM mask excluded) as protocols were added in the optimal whole-WM order (ADRC, MESO-72, UK Biobank, ADNI, MESO-96, ABCD, HCP-DA1), then averaged across regions. Each value uses the noise-corrected estimator, applied to that region's fitted protocol means and their covariance, and is expressed relative to the region's intercept, $\bar{Y}$.**(A)** Pooled $CV_{protocol}$ at each step, computed as the square root of the sum of the squared region-averaged CVs of the nine metrics. Bar segments show each metric's share of that sum.**(B)** Region-averaged $CV_{protocol}$ for each metric at each step, by family (DTI, DKI, SMI). Shaded bands are 95% confidence intervals from subject resampling (200 replicates). In each replicate, $CV_{protocol}$ was recomputed in every region and averaged across regions; the bands span the 2.5th to 97.5th percentiles of these averages.

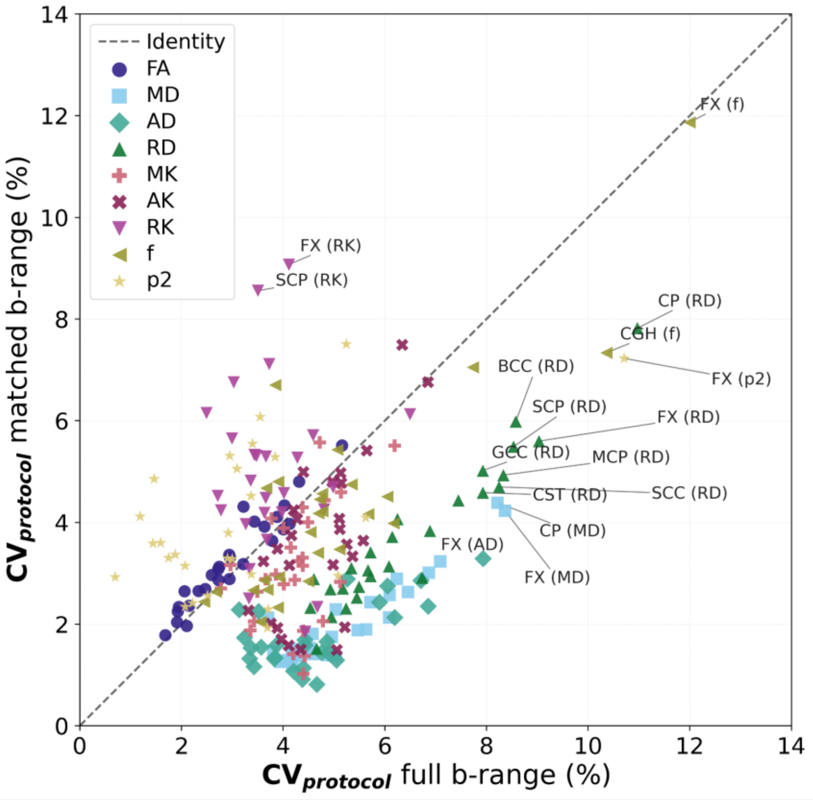


**Supplementary Figure S7**. *Effect of matching the b-value range on inter-protocol variation***.** Each point is $CV_{protocol}$ for one white matter region and one metric, with the full b-range on the x-axis and a *b*-range matched across protocols on the y-axis. The full *b*-range uses all acquired shells. The matched *b*-range restricts DTI estimations to b ≤ 1000 s/mm² and DKI/SMI estimations to b ≤ 2000 s/mm². Because HCP-DA1 includes higher shells, the matched analysis uses HCP-DA2b instead. Both axes use the same denominator, the average of the two analyses' intercepts, so distance from the identity line (dashed) reflects only the change in spread between protocols. Points below the line have lower variation between protocols when *b*-ranges are matched. Values are averaged across hemispheres, and the whole white matter mask is excluded. Labeled points are outliers, either with CV above Q3 + 1.5 × IQR or with a full versus matched difference beyond 3 × IQR.

**Registration Bias Testing**

We explore diffeomorphic atlas registration itself as a potential source of variability and bias. First, we examined the log-Jacobian determinant of the atlas-to-native registrations across protocols. Second, we compared $CV_{protocol}$ computed in each acquisition's native space with $CV_{protocol}$ computed after transformation to a subject-specific template. In short, the deformation applied by the registration reflects mainly anatomical differences between subjects rather than the acquisition protocol, and the small protocol dependent component has no meaningful effect on the reported inter-protocol variability.

We examined the log-Jacobian determinant (log *J*) of each atlas-to-native warp, across the whole brain and within the JHU regions used in our analyses (**Supplementary Figure S8**). Across the whole brain, the distribution of log *J* had the same shape and width for every protocol (**Supplementary Figure S8A**). The amount of deformation, measured as the within-map SD of log *J*, differed between protocols by no more than it differed between repeat scans of the same protocol. Protocols differed mainly by a small shift of the whole distribution, with mean log *J* ranging from −0.009 (HCP-DA1) to +0.070 (ADRC). Within the 27 JHU regions, subject anatomy accounted for most of the variance in regional log *J* (median 81% across regions), and protocol for a smaller share (median 8%; **Supplementary Figure S8B**). Registration therefore deforms each protocol by a similar amount, with a modest protocol-dependent shift.

Second, we tested whether this protocol-dependent shift affects our results by repeating the analysis in a subject space. In this space, all protocols from a subject share a single nonlinear warp through a study-specific template and differ only by a rigid transformation, which removes any protocol-dependent deformation. $CV_{protocol}$ changed little (**Supplementary Figure S9**). For seven of nine metrics the two spaces were statistically equivalent within ±2 percentage points, and the median absolute difference across regions was 0.2 to 0.7 percentage points. The exceptions, MK and AK, reflected unstable subject-space fits in the brainstem and corticospinal tract. Native space also performed better. It gave lower $CV_{protocol}$ for FA, MD, AD, and RD, and similar values for the other metrics. All model fits that failed did so in subject space. Removing protocol-specific warps therefore did not reduce inter-protocol variability, and registration does not drive the reported results. We kept the native-space approach because it is more robust, avoids resampling the diffusion data, and does not require repeated scans of each subject to build a subject template. That makes it applicable to large cross-sectional studies.

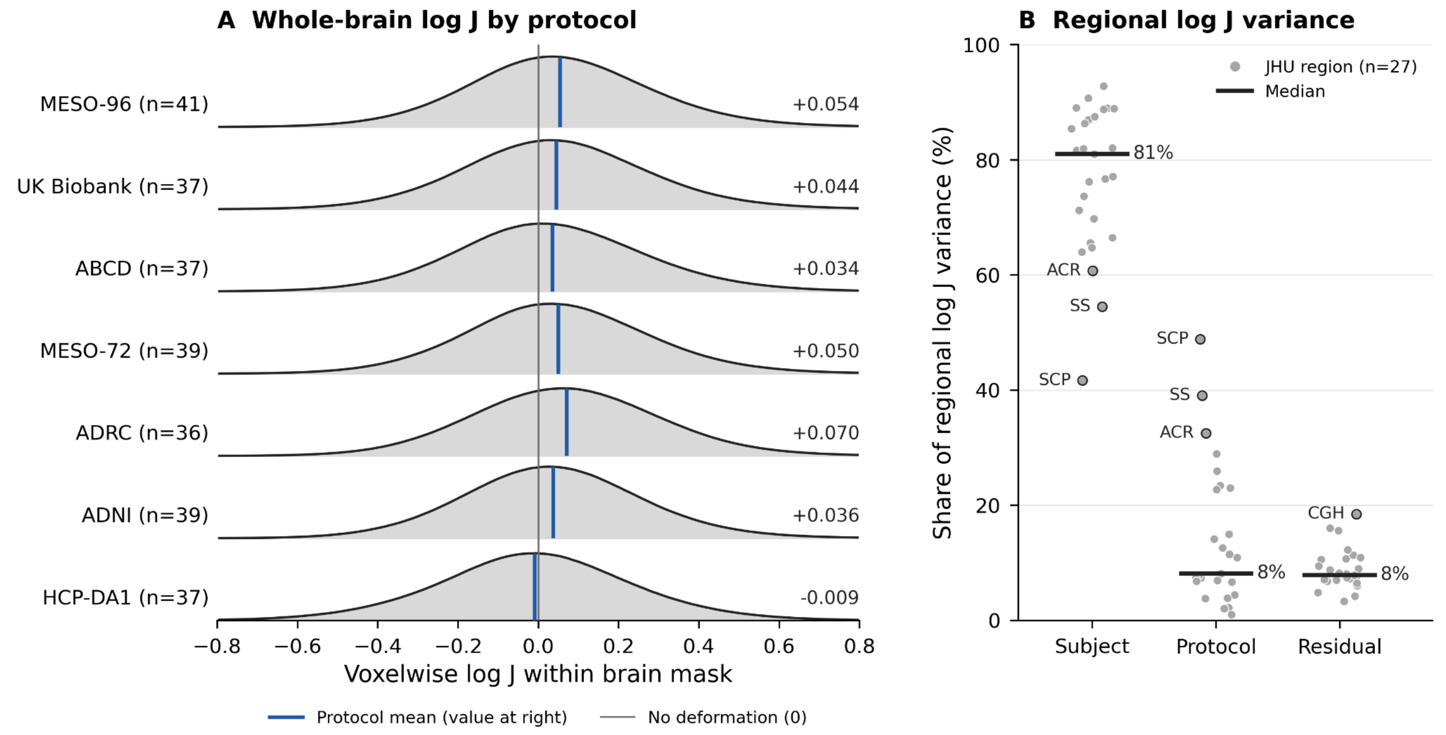


**Supplementary Figure S8.** *Log-Jacobian of the atlas-to-native registrations.* The JHU-ICBM FA atlas (1 mm) was registered to each acquisition's native FA map with ANTs (rigid, affine, then SyN), with the atlas as the fixed image. The log-Jacobian determinant (log $J$) of the nonlinear (SyN) warp was computed on the atlas grid. Because the affine component is excluded, log $J$ measures local deformation after global size and shape are removed. Log $J > 0$ indicates local expansion of the subject relative to the atlas, and log $J < 0$ indicates contraction. Data are from the seven main protocols, restricted to acquisitions passing SQUAD quality control and the registration check (fraction of warped atlas labels inside the brain mask ≥ 0.95; all acquisitions passed). The two MESO-96 runs were averaged within each subject. **(A)** Distribution of voxelwise log $J$ within the atlas brain mask for each protocol. Each scan's histogram (bin width 0.01) was normalized to a density, averaged over the MESO-96 runs, and then averaged over subjects; $n$ is the number of subjects. Blue lines mark each protocol's mean log $J$ (value at right), and the gray line marks no deformation (log $J = 0$). **(B)** Share of the variance in regional log $J$ attributable to subjects, protocols, and residual error. The shares come from the LME of protocol variance model, fitted separately in each of 27 JHU regions (left and right hemispheres averaged, whole white matter mask excluded) with the regional mean log $J$ as the outcome. The protocol share uses the noise-corrected spread of the fitted protocol means. Gray points are regions and black bars are medians across regions. Labeled regions are the three with the largest protocol share, the superior cerebellar peduncle (SCP), sagittal stratum (SS), and anterior corona radiata (ACR), shown in the subject and protocol columns, and the one region beyond 1.5 × IQR of the residual share, the hippocampal cingulum (CGH).

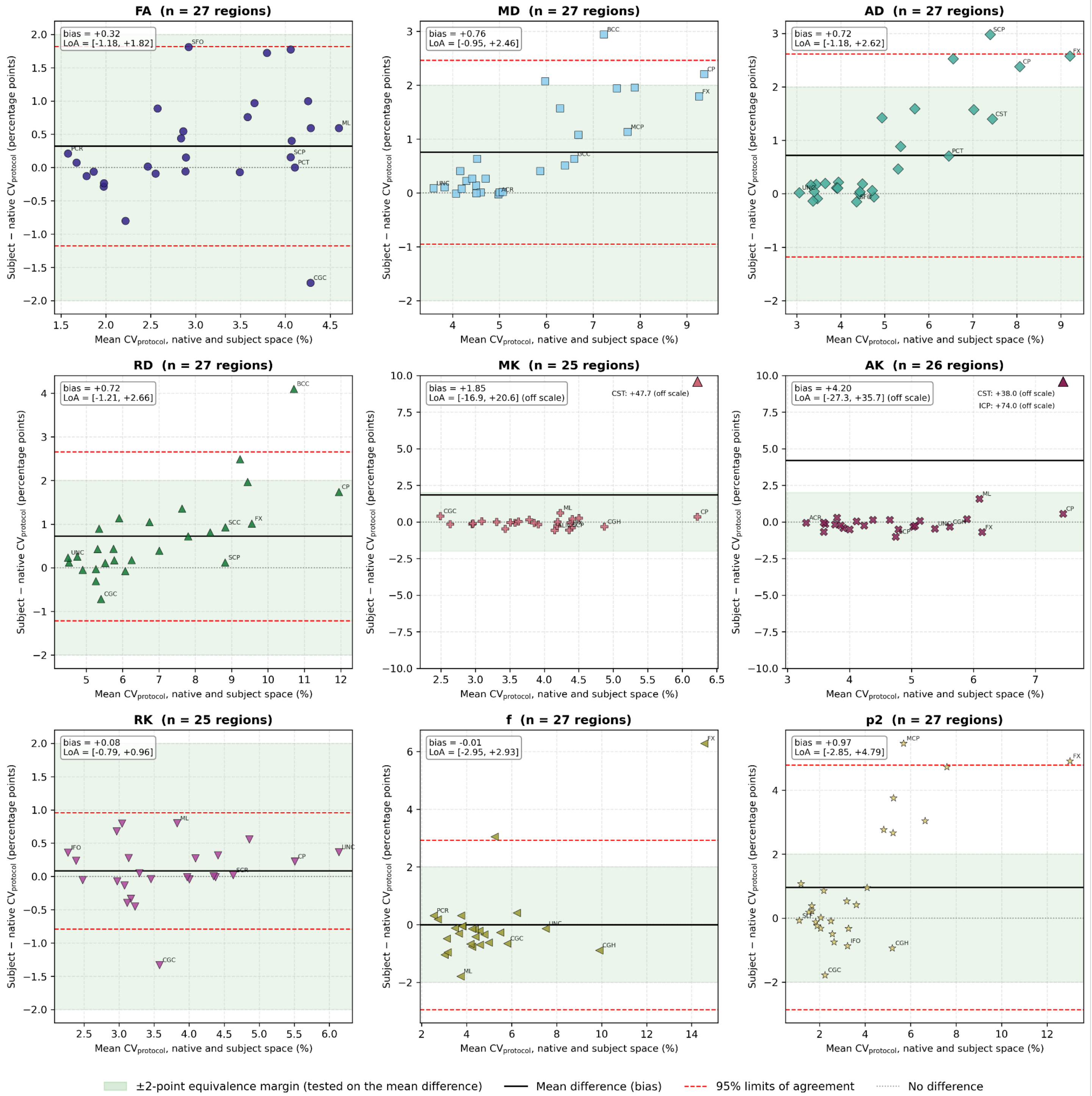


**Supplementary Figure S9.** *Agreement of $CV_{protocol}$ between native-space and subject template-space registration*. Bland-Altman plots (Bland & Altman, 1986) comparing $CV_{protocol}$ computed with two strategies for placing the JHU atlas labels. In "native" space, used throughout the manuscript, the atlas was warped directly to each acquisition with a single nonlinear (SyN) registration, and the diffusion maps were not resampled. In "subject" (subject template) space, the labels reached each subject through study-specific templates (atlas to cohort template to subject template), and each acquisition's diffusion maps were aligned to the subject template by a rigid transformation only, so all protocols from a subject share one nonlinear warp. In each space, $CV_{protocol}$ was estimated from the

LME of the protocol variance model, fit separately for each metric and region using the mean ROI values of the seven main protocols. Left and right hemispheres were averaged and the whole white matter mask was excluded, giving the 27 regions. Only subject × protocol × region combinations present in both spaces and passing SQUAD quality control were included. Five cells were excluded because the subject-space fit was singular: MK, AK, and RK in the middle cerebellar peduncle, and MK and RK in the inferior cerebellar peduncle. Each point is one region ($n$ = 27; 25 for MK and RK, 26 for AK). The x-axis shows the mean of the two $CV_{protocol}$ values and the y-axis the difference, subject minus native, in percentage points. The solid line is the mean difference (bias), the dashed red lines are the 95% limits of agreement (bias ± 1.96 SD), and the dotted line marks no difference. The shaded band is the ±2-percentage-point equivalence margin, tested on the mean difference (two one-sided tests, FDR-corrected); the two spaces were equivalent for all metrics except MK and AK. Regions beyond the plotted range are shown as a triangle at the upper edge with their differences listed (MK: CST; AK: ICP and CST). These reflect unstable subject-space fits and account for the wide MK and AK limits of agreement. In each panel the most discrepant regions and those at the extremes of either axis are labeled.
